# Incorporating Multi-threshold Polygenic Risk in Hippocampal-based Normative Models Improves Cognitive Decline Prediction

**DOI:** 10.1101/2025.04.03.646991

**Authors:** Mohammed Janahi, Luigi Lorenzini, Neil P. Oxtoby, Frederik Barkhof, Younes Mokrab, Johnathan M Schott, Andre Altmann, the Alzheimer’s Disease Neuroimaging Initiative

**Author notes:** Data used in the preparation of this article were obtained from the Alzheimer’s Disease Neuroimaging Initiative (ADNI) database (adni.loni.usc.edu). As such, the investigators within the ADNI contributed to the design and implementation of ADNI and/or provided data but did not participate in analysis or writing of this report. A complete listing of ADNI investigators can be found at: http://adni.loni.usc.edu/wp-content/uploads/how_to_apply/ADNI_Acknowledgement_List.pdf.

## Abstract

Predicting cognitive decline using MRI measures of brain volume is challenging. Normative models capture the evolution of biomarkers over time in a healthy group, quantifying an individual’s deviation from that norm. Here, we improve the precision of normative models for hippocampal volume, a key cognitive biomarker, by integrating multi-threshold polygenic scores (PGS) that capture genetic predisposition to its variability. We built Gaussian Process Regression (GPR) models on genetic, imaging, and demographic data from 23,997 UK Biobank (UKBB) participants, validating on 3,000 out-of-sample participants from the Alzheimer’s Disease Neuroimaging Initiative (ADNI) and the European Prevention of Alzheimer’s Disease (EPAD) cohorts. The genetically-informed models improved associations significantly across five experimental designs and 13 key neurocognitive measures, including Mini-Mental State Examination (MMSE), Clinical Dementia Rating (CDR), Alzheimer’s Disease Assessment Scale (ADAS), and importantly, improving prediction of future cognitive decline. These findings underscore the value of integrating multi-threshold PGS with neuroimaging-based predictive models for improving prognostication in neurodegenerative diseases.

## 1. Introduction

Brain imaging provides invaluable insights into the pathophysiology of brain disorders. Over the past decades, various case-control studies have identified multiple disease hot-spots in the brain, notably large-scale meta-analyses by the ENIGMA consortium which established prototypical abnormality maps for dozens of brain disorders [1]. However, the paradigm of case-control studies assumes that all cases express the same disease-related imaging pattern, which is unlikely given the heterogenous nature of most brain disorders.

Increasingly, normative modelling is used in neuroimaging research, comparing individual subjects’ brains to the expected appearance in a healthy cohort to identify personalized disease maps [2–4]. As such, normative modelling was shown to be effective in modelling heterogeneous diseases like schizophrenia and attention deficit hyperactivity disorder (ADHD) [2, 5]. Recent work has established normative brain models mapping the entire lifespan based on over 100,000 participants [6]. Janahi et al. demonstrated that such models can be improved by incorporating polygenic scores (PGS), which capture genetic predisposition to variation in a brain phenotype of interest such as hippocampal volume (HV), thereby allowing abnormality detection up to three years earlier [7]. Other studies have similarly found significant benefits of genetic adjustment in normative modelling [8, 9].

PGS are typically constructed as a weighted combination of Single Nucleotide Polymorphisms (SNPs) [10], deriving the weights from Genome Wide Association Studies (GWAS)[11]. PGS were calculated to measure genetic risk across a range of clinical phenotypes including coronary artery disease, diabetes, and Alzheimer’s disease (AD) [12]. Furthermore, PGSs were used to estimate long term health trajectories in Schizophrenia, AD and ADHD [13–17].

Linkage Disequilibrium (LD) poses a challenge when creating PGSs. While GWAS estimates independent SNP effects, SNPs are often inherited not individually, but as haplotype blocks; [18] consequently, PGS construction needs to adjust for LD. The standard method for LD adjustment is Clumping and Thresholding (C+T), where SNPs below a certain p-value threshold are selected, followed by clumping neighbouring SNPs from the same LD haplotype, then retaining the highest effect size SNP in each clump [19]. For the remaining SNPs, the effect sizes are then weighted by an individual’s genotype (0,1, or 2 copies of the SNP) and summed to produce the PGS. More advanced methods adjust all GWAS SNP effect estimates, combining all SNPs without clumping or thresholding, assuming either a small (e.g., LD-Pred [20], SBayesR [21]) or a large number of diseasing-causing SNPs (e.g., PRS-CS [22], lassosum [23], SBLUP [24]). However, C+T method remains the most widely used [25] assuming a wide range of P-value thresholds ranging from stringent (P<5e-8) and selecting SNPs with low standard errors, to lenient cut-offs (P<0.5) allowing better coverage but less accurately estimated effects. While a single best performing threshold is often selected for downstream analysis, there is mounting evidence that PGSs at different thresholds capture different aspects of the trait and are not simply interchangeable[26–29]. For instance, Stacked Clumping and Thresholding which combines approximately 100,000 PGS using a penalized logistic regression, leads to improved disease prediction performance[30].

In this study, we explore the value of combining multiple P-value thresholds to improve normative models for HV and its clinical application for prognosis of cognitive decline. First, we assess the information in HV PGSs at different thresholds and use machine learning approaches to leverage them (Figure 1A). Next, we integrate the combined PGSs into multivariate normative models based on data from UK Biobank (UKBB) [31] and evaluate them in two out-of-sample cohorts across the AD spectrum: the Alzheimer’s Disease Neuroimaging Initiative (ADNI) database [32], which comprises cognitively normal subjects and subjects with mild cognitive impairment and AD, and the European Prevention of Alzheimer’s Disease (EPAD) database [33], which comprises cognitively normal participants at risk for dementia (Figure 1B). Finally, we compare the models to standard in terms of correlations to various neurocognitive measures, including memory and orientation, both cross-sectionally and longitudinally.

**Fig. 1:**
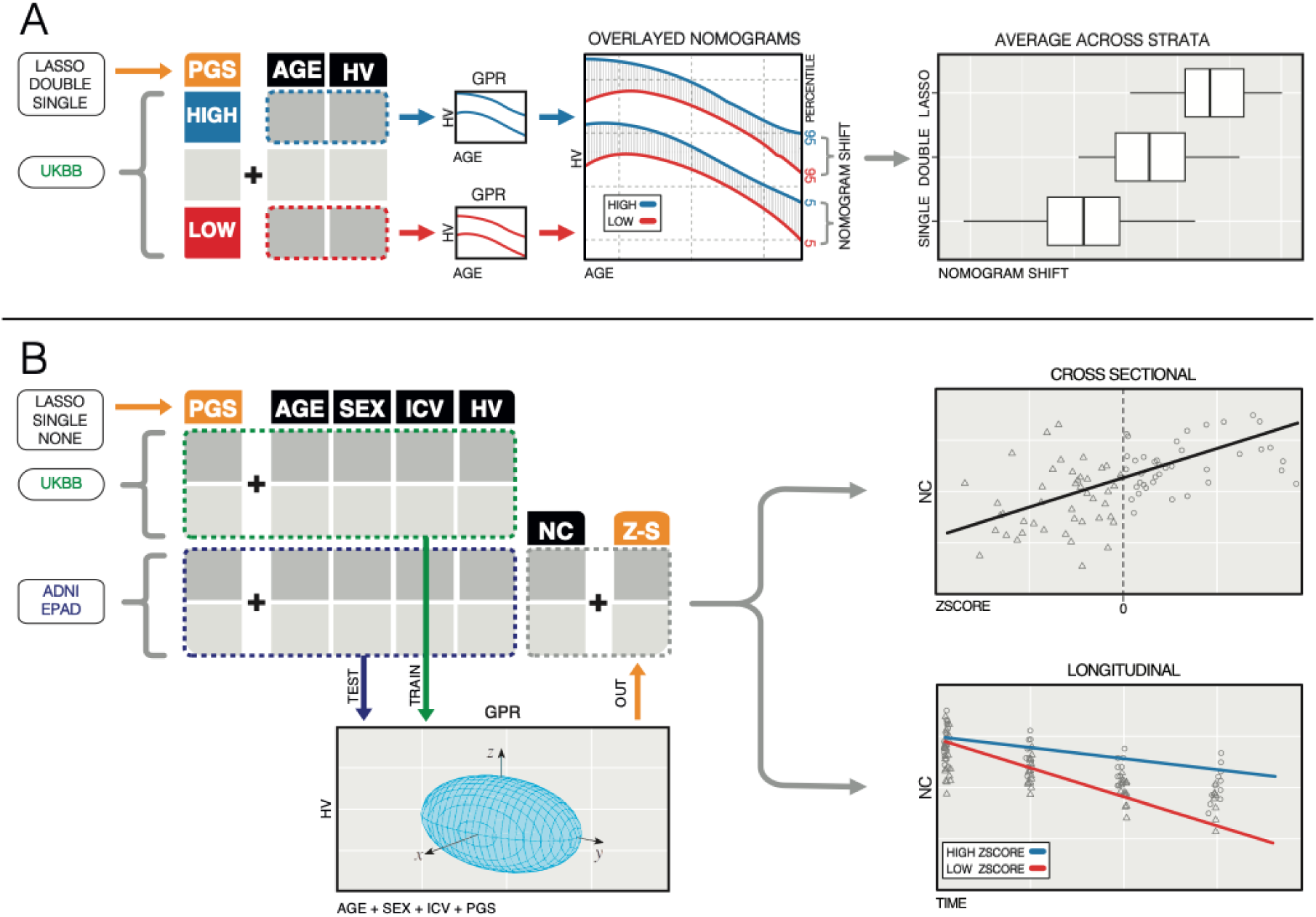
Study Overview. To explore the value of combing multiple p-value thresholds of the same C+T based Polygenic Scores (PGS) in predicting cognitive decline, (A) we generate PGS using single, pairs, and 13 combined thresholds (LASSO-PGS) for UK Biobank (UKBB). We then train gaussian process regression (GPR) models on high/low PGS participants and report the predicted model differences. (B) Next, we train UKBB models with no PGS, single threshold PGS, or LASSO-PGS and test them in two out-of-sample, harmonized datasets (ADNI, EPAD) to obtain HV z-scores. We finally correlate the z-scores to multiple Neurocognitive measures such as memory and orientation cross-sectionally and longitudinally.

## 2. Results

### 2.1 PGS at different thresholds contain complementary information

We built normative models using data from UKBB participants. In total we included data from 23,997 participants, after applying exclusion criteria for existing health conditions and genetic ancestry (Table 1). To measure the impact of the genetic information, participants were stratified into two groups: genetically predicted larger and smaller hippocampi.

**Table 1:** Summary of Pre-processed Datasets.

|  | UKBB (n=23,997) | EPAD (n=1,755) | ADNI (n=1,245) |  |  |
| --- | --- | --- | --- | --- | --- |
|  |  |  | CN (n=479) | MCI (n=617) | AD (n=149) |
| Age (mean, SD) | 64.4 (7.5) | 65.8 (7.1) | 71.5 (5.2) | 71.0 (6.7) | 71.6 (7.2) |
| Sex |  |  |  |  |  |
| Male | 11799 (49.2%) | 774 (44.1%) | 198 (41.3%) | 356 (57.7%) | 84 (56.4%) |
| Female | 12198 (50.8%) | 981 (55.9%) | 281 (58.7%) | 261 (42.3%) | 65 (43.6%) |
| HV mm3 (mean/SD) | 4043 (398) | 3994 (408) | 3938 (339) | 3679 (401) | 3418 (399) |
| MMSE (mean/SD) |  | 28.4 (1.8) | 29.0 (1.1) | 27.9 (1.7) | 23.2 (2.1) |
| CDR-SB (mean/SD) |  | 0.36 (0.74) | 0.03 (0.13) | 1.4 (0.8) | 4.3 (1.5) |

HV: Hippocampal Volume (after cross-dataset harmonization; Methods), MMSE: Mini-Mental State Exam, CDR-SB: Clinical Dementia Rating

Following previous work by Janahi et al [7], separate nomograms were estimated for both strata and the degree of separation was computed (Methods). PGS were generated using PRSice-2 [34] on harmonized SNP data (Methods). The most lenient (P=1) and most stringent (P=1E-08) P-value cutoff comprised 70,251 and 4 SNPs, respectively.

#### 2.1.1 SINGLE THRESHOLD PGS CORRELATIONS

HV PGSs at different thresholds exhibited consistent correlations with measured HV, with correlations being strongest at the opposing ends (Figure 2 A, Supp. Figure 1). PGSs using similar P-value thresholds show strong correlations between each other, while PGSs using distant P-value thresholds appear uncorrelated (e.g., P=1E-8 and P=1.0 exhibit Pearsons *r* =.06) (Figure 2 C, Supp. Figure 2).

**Fig. 2:**
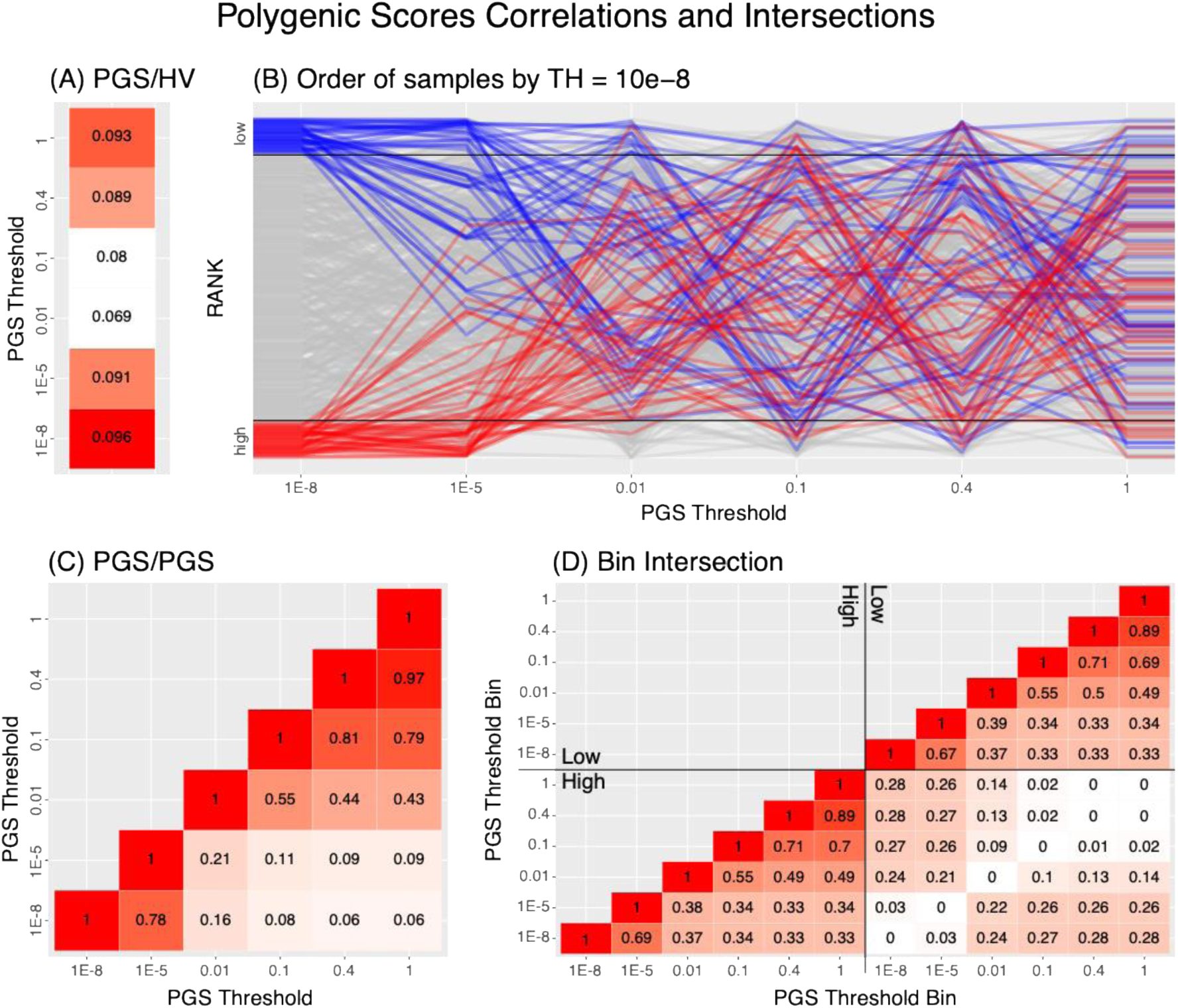
HV PGS Correlations in UKBB. (A) HV PGSs generated at different P-value thresholds correlate similarly to HV (P=1E-8 & P=1 exhibit Pearsons *r* = .09). (B) The order of participants is not preserved when ranking them by the P=1E-8 threshold PGS. The top and bottom 10% of participants (blue/red lines) are essentially randomly distributed among the ranks of participants by the P=1 threshold PGS. (C) These same PGS become less correlated as the distance between the thresholds increases; ranging from r=.97 down to r=.06 (D) Overlap of participants ranked in the top/bottom 30% by each PGS. Participants ranked high/low by two polygenic scores (bottom-left/top-right) overlap more than those ranked high by one score but low by the other (bottom-right). For instance, 33% of the participants ranked as high at the P=1E-8 threshold are also ranked high at the P=1 threshold. However, 28% of the participants ranked high at the P=1E-8 threshold are also ranked low at the P=1 threshold. The diagonal of the bottom right sector of the table is all zeros since it represents the intersections of the top and bottom bins in the same threshold PGS.

When considering an integration of PGS into normative models, it is important to consider the ranking the PGS generates for the participants. Indeed, when subjects are ranked with respect to a PGS using P-value threshold 1E-8, then they appear nearly randomly distributed when viewed at the P-value threshold of P=1.0 (Figure 2B, Supp. Figure 4). That is, PGSs at different thresholds split participants differently. For instance, of the subjects ranked high according to the P=1E-8 threshold (i.e., highest 30% of subjects by HV PGS), only 33% were also ranking high using the P=1 threshold, while 28% were ranked low at the P=1 threshold and the remaining 39% were ranked In the middle (Figure 2D; Supp. Figure 2/3). As before, overlap between groups was higher when the two P-value cutoffs were closer.

#### 2.1.2 LASSO-PGS

These previous analyses showed that the HV-PGSs constructed at different threshold carry complementary information. Next, we used a LASSO regression [35] to combine the scores at different cutoffs into single score (LASSO-PGS), which we hypothesize to better correlate with HV than single scores (Methods). To this end we trained 30 bootstrap replicates of sex-stratified models on UKBB. The LASSO model consistently put higher weights on the scores with the highest and lowest thresholds (Supp. Figure 5).

### 2.2 Combining PGSs improves Nomogram Separation

To evaluate the effect of the different PGS on the normative models, we estimated different nomograms for the subjects ranked in the highest and lowest 30% according to the PGS. Next, we computed the level of Nomogram Separation between the “high” and the “low” groups (Methods). This was conducted for the LASSO-PGS and as a reference for all single PGS thresholds and all combinations of two thresholds (double-PGS). Compared to a single PGS threshold, using the double- or LASSO-PGS improved Nomogram Separation. On average, double threshold PGSs were 22% better than single threshold PGS, and that the LASSO-PGS was 50% better (Figure 3, Supp. Figure 6/7).

**Fig. 3:**
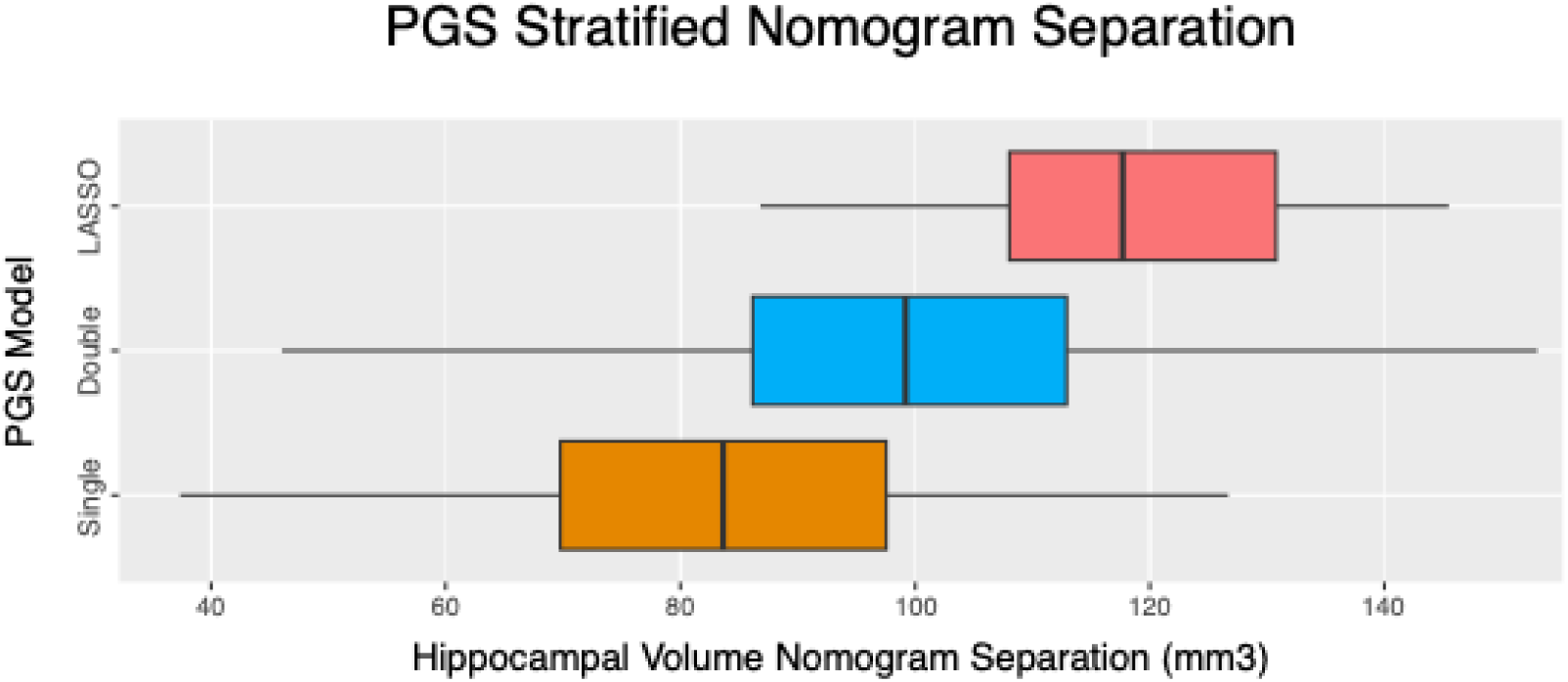
Combining multiple PGS improves Nomogram Separation. Comparing the nomogram separation in UKBB across 30 bootstraps of stratification by 13 single threshold PGSs of HV; 78 combinations of two thresholds; and a lasso of all 13 thresholds. On average, a double-PGS is 22% better than a (classic) single threshold PGS, and a LASSO-PGS is 50% better.

### 2.3 Combining PGSs improves HV models in out-of-sample datasets

After having established that combining multiple PGS using a LASSO regression provides a better separation between nomogram strata, we aimed to build multivariate normative models that can be applied to external datasets to improve diagnosis and prognosis of people with dementia. After applying inclusion criteria based on genetic and imaging data availability and matching genetic ancestry, 23,997 participants were included from UKBB for training, and for evaluation 1,245 participants (6,814 visits) from ADNI and 1,755 participants (4,938 visits) from EPAD (Table 1).

#### 2.3.1 Harmonization and Cross Dataset Common Polygenic Score

To have comparable PRS across datasets, the SNP content across all three databases was harmonized (resulting in 262,660 common SNPs; Methods) and PRS were recomputed at the 13 cutoffs with the most lenient cutoff (P=1.0) comprising 53,686 SNPs. Dataset specific biases were removed prior to further downstream analysis (Methods; Supp. Figure 8).

#### 2.3.2 Model Training

We randomly split UKBB into training (50%) and testing (50%) cohorts and trained different normative models using Gaussian Process Regression (GPR). For all models, HV was the target variable, and all models contained age (A), sex (S) and intracranial (I) volume as predictors (“ASI” models). In addition, we trained models that used a single PRS as predictor (ASIP-single) and models that used the LASSO-PRS as predictor (ASIP-LASSO). The trained models were applied to the UKBB test cohort, and to both ADNI and EPAD participants. We found that GPR models improved when incorporating genetic information, with the strongest improvement being seen with the LASSO-PGS (Figure 4). Unsurprisingly, RMSE increased when going from training to testing in UKBB and further when applying the models to out-of-sample datasets. When evaluating the model performance by diagnosis, EPAD and CN participants in ADNI exhibited similar performance compared to participants in the UKBB test set (Supp. Figure 10). Thus, indicating that larger deviations in performance are caused by disease effects, demonstrating the underlying assumption of normative models. Overall, the relative performance of all tested strata improved when adding single threshold PGS and further with LASSO-PGS.

**Fig. 4:**
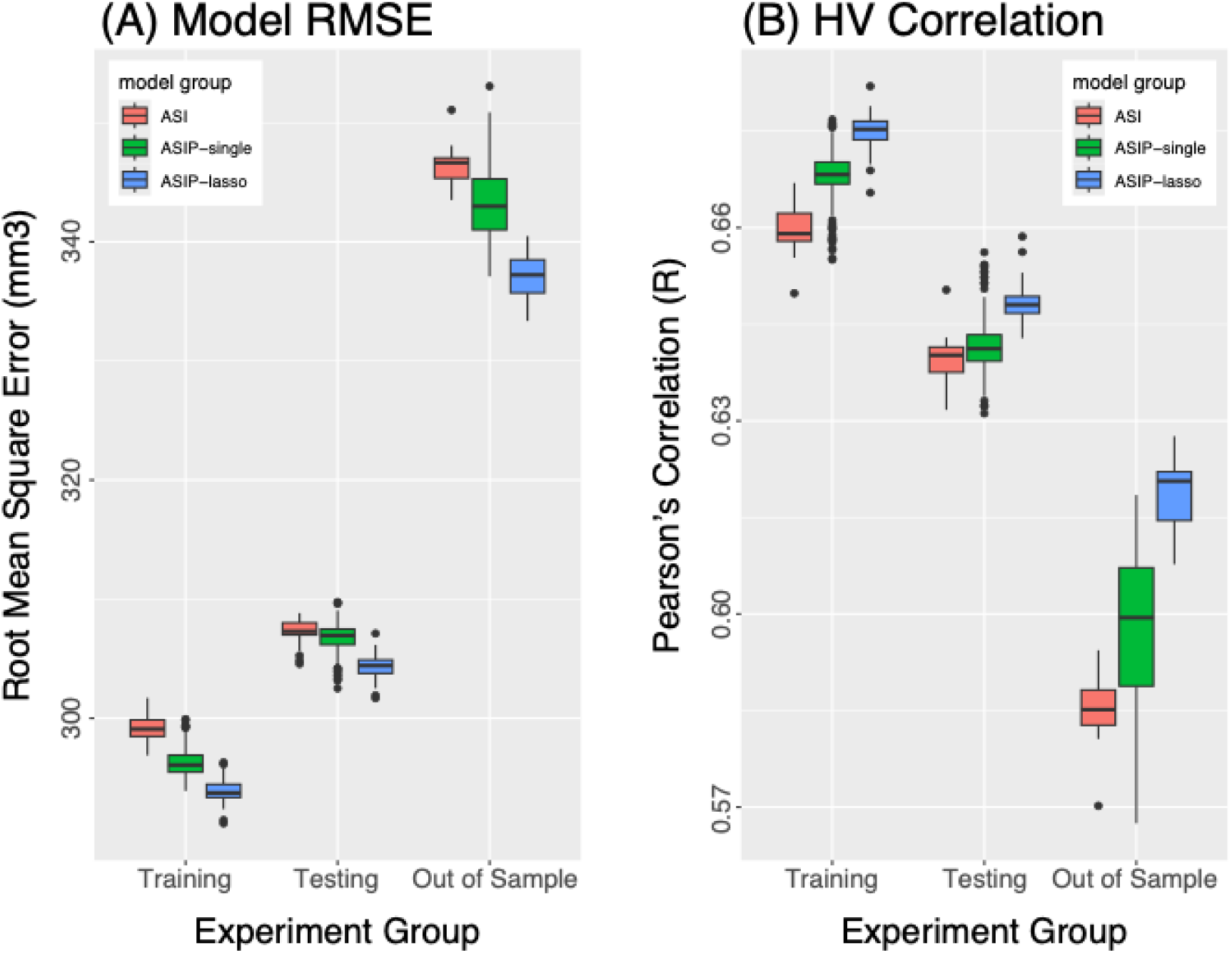
High Dimensional GPR models across datasets and covariates. (A) Root Mean Squared Error (RMSE) between predicted HV by the GPR models and measured HV. On average, the RMSE lowers when going from the standard Age-Sex-Intracranial (ASI) model (red) to the ASI + polygenic (ASIP) using single threshold PGS (ASIP-single; green), to the ASIP-LASSO model (blue). RMSE increases on test data and out-of-sample cohorts. (B) Pearson’s correlation coefficient between predicted and measured HV by the different models.

### 2.4 Combining PGSs Improves Prediction of Cognitive Performance Cross-Sectionally And Longitudinally

In a final set of experiments, we challenged the clinical utility of these genetically adjusted normative models. We used the rich phenotyping data available in both ADNI and EPAD to assess whether adding genetics into the normative models improves the association with current and future cognitive function. In these experiments we computed participants’ percentile score using the ASI model as well as the ASIP-LASSO model. Next, using linear models and linear mixed effect models we tested whether the percentile shift induced by the ASIP-LASSO model compared to the ASI model (termed *model-shift*) better explains current and future test scores, respectively (Methods). Cross-sectionally, we found significant correlations to multiple neurocognitive (NC) tests and domains across both out-of-sample datasets. In both ADNI and EPAD, the model-shift introduced by the LASSO-PGS demonstrated significant added benefit over the classic normative model (ASI) for MMSE and CDR-SB (Figure 5, Supp. Figure 11/12, Table 2, Supp. Table 1). In ADNI, ASIP-LASSO significantly improved the association with ADAS13 and MOCA. When limiting the analysis to participants showing moderate model-shift (<11%), there was significant association with CDR-SB and Flanker in ADNI and EPAD, respectively. Results were similar when comparing ASIP-LASSO to ASIP-single; even after excluding participants with dementia from ADNI (Supp. Table 1). Model-shift was significantly associated to all cognitive domains in ADNI. These associations remained significant when only participants with moderate shift (<11%) were considered, i.e., these results were not driven by few extreme cases with strong model shift. In EPAD, the model-shift was significantly associated with Memory and Executive Function (Table 2).

**Fig. 5:**
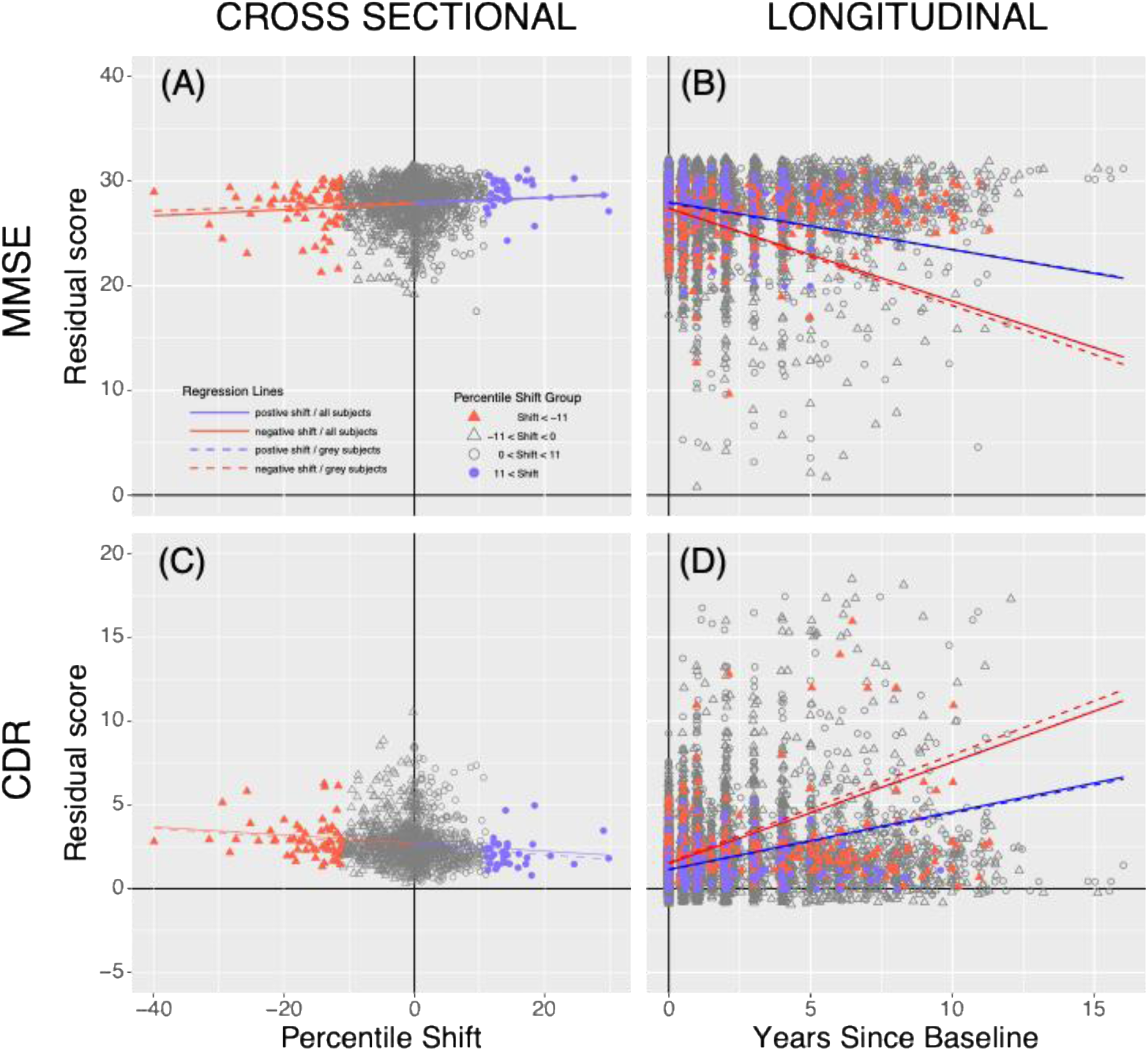
Cross-sectional and Longitudinal Effect of Model-shift. Illustrative example of comparing performance in MMSE and CDR-SB in ADNI to ASIP-LASSO to ASI model shift. (A, C) At baseline, model shift is positively correlated to MMSE (P=2.6E-3), where a higher score is better; and negatively correlated to CDR (P=7.7E-5), where a lower score is better. When observing moderately shifted participants (|model-shift|<11%; grey points) the CDR-SB correlation remains significant (P=3.6E-3) and the magnitude are not significantly changed (dashed line). (B, D) These correlations remain significant longitudinally (MMSE: P=3.6E-4, CDR-SB: P=8.4E-4). The regression lines drawn are projections/prediction for an average subject with +10 vs −10 model-shift (blue vs red).

**Table 2:** Neurocognitive test/domain association with model-shift.

| TABLE | NC Test /<br>Domain | Cross Sectional Results |  |  |  | Longitudinal Results |  |  |  |
| --- | --- | --- | --- | --- | --- | --- | --- | --- | --- |
|  |  | All |  | No High Shift |  | All |  | No High Shift |  |
|  |  | slope | p-val | slope | p-val | slope | p-val | slope | p-val |
| ADNI | MMSE | .029 | **2.6E-03** | .021 | 1.3E-1 | .022 | **3.6E-04** | .025 | **5.4E-03** |
|  | CDR-SB | -.024 | **7.7E-05** | -.026 | **3.6E-3** | -.013 | **8.4E-04** | -.015 | 7.5E-03 |
|  | ECOG.SP | -.008 | 1.0E-02 | -.008 | 7.6E-2 | -.0009 | 2.1E-01 | -.001 | 3.3E-01 |
|  | ADAS13 | -.149 | **2.1E-05** | -.129 | 1.1E-2 | -.033 | 1.1E-02 | -.044 | 2.2E-02 |
|  | MOCA | .053 | **5.5E-03** | .059 | 2.6E-2 | .012 | 1.2E-02 | .013 | 7.4E-02 |
|  | Memory |  | **5.8E-10** |  | **9.4E-04** |  | 2.4E-01 |  | 1.6E-01 |
|  | Language |  | **1.8E-03** |  | **9.4E-04** |  | **5.9E-03** |  | 3.5E-02 |
|  | Orientation |  | **1.4E-04** |  | **2.1E-02** |  | **4.5E-06** |  | **5.5E-04** |
|  | Visuospatial |  | **1.5E-02** |  | **2.1E-02** |  | 1.5E-01 |  | 7.0E-01 |
|  | Exec. Function |  | **2.5E-05** |  | **3.4E-03** |  | **1.4E-03** |  | 8.0E-02 |
| EPAD | MMSE | .019 | **3.0E-03** | .014 | 1.6E-1 | .004 | 8.8E-01 | .002 | 8.0E-01 |
|  | CDR - SB | -.007 | **9.2E-03** | -.008 | 3.8E-2 | -.0005 | 8.5E-01 | -.001 | 4.7E-01 |
|  | RBANS | .112 | 3.9E-02 | .139 | 9.2E-2 | -.005 | 6.1E-01 | .025 | 5.7E-01 |
|  | Flanker | .007 | 5.8E-02 | .015 | **6.5E-3** | -.002 | 1.3E-01 | -.003 | 2.5E-01 |
|  | Dot Count | .033 | 1.0E-01 | -.005 | 8.7E-1 | -.002 | 1.2E-01 | .035 | 1.7E-01 |
|  | Memory |  | **9.6E-05** |  | **4.0E-03** |  | 8.9E-01 |  | 9.4E-01 |
|  | Language |  | 6.9E-01 |  | 4.3E-01 |  | 4.8E-01 |  | 2.6E-01 |
|  | Orientation |  | 1.8E-01 |  | 2.5E-01 |  | 1.9E-01 |  | 4.3E-01 |
|  | Visuospatial |  | 8.1E-02 |  | 2.6E-01 |  | 5.7E-01 |  | 1.5E-01 |
|  | Exec. Function |  | **4.0E-05** |  | 5.3E-02 |  | 1.7E-01 |  | 3.4E-01 |

Note: Bold items: statistically significant after Holm–Bonferroni correction across 5 sets of tests/domains in each dataset independently. Correction was done independently in cross-sectional/longitudinal, and all/no-high-shift groups. Cross sectional P-values are derived from linear models, and longitudinal p-values are derived from an ANOVA between two Linear Mixed Effect (LME) models (see Methods). MMSE: Mini-Mental State Examination; CDR-SB: Clinical Dementia Rating Sum of Boxes; ECOG SP: Everyday Cognition Study Partner; ADAS13: AD Assessment Scale13 items; MOCA: Montreal Cognitive Assessment; RBANS: Repeatable Battery for the Assessment of Neuropsychological Status

In our longitudinal linear mixed effects models, model-shift was significantly associated with MMSE and CDR-SB in ADNI (Figure 5, Table 2, Supp. Table 1), with shifts towards larger percentiles indicating a longitudinally better cognitive profile. In the domain tests, we found significant longitudinal association with Language, Orientation, and Executive Function (P=5.9E-3, P=1.5E-5, P=1.4E-3, respectively) (Table 2). There were no significant longitudinal associations in the EPAD cohort. Likely owing to the short-term follow-up data compared to ADNI.

Finally, percentile shifts are not uniformly meaningful; crossing the 50^th^ percentile may have less significance than crossing the 5^th^ percentile for example. To test whether this was the case in our models, we recorded whether the ASI to ASIP-LASSO *model-shift* crossed predefined percentile thresholds and tested whether these crossings were associated with longitudinal performance using linear mixed-effects models (Methods). In ADNI, crossing at the 1st and 10th percentile thresholds was significantly associated with performance in multiple neurocognitive tests and all domains (Supp. Figures 13 and 14, Supp. Tables 1 and 4). For example, MMSE and CDR-SB showed significant effects at the 1^st^ percentile (P= 4.3E-15 and P=2.1E-9) and the 10^th^ percentile (P=4.5E-7 and P=2.3E-7).

## 3. Discussion

In this study, we hypothesised that PGS constructed at different thresholds capture different spectra of polygenic traits and that combining multiple threshold PGSs of the same phenotype will improve their predictive performance. Our experiments, focused on hippocampal volume (HV), a key biomarker in Alzheimer’s disease, confirmed that different PGS thresholds contain distinct information about HV. While all PGS correlated well with HV, they exhibited different degrees of correlation amongst themselves (ranging from *R*=0.06 to *R*=0.97; Figure 2-C). Furthermore, when participants were ranked based on a PGS at one cutoff (e.g., 1E-8), their ranks appeared essentially random when viewed at another PGS cutoff (Figure 2-B). This result confirms previous observations in PGS threshold studies [30, 36, 37]. Consequently, the proposed LASSO-PGS models were able to leverage the complementary information and improved nomogram separation, model fit, and predictions of neurocognitive test performance compared to models not using any polygenic information (no-PGS) or classic single-threshold-PGS models. In Nomogram Separation Experiments (Figures 2 and 3) we performed tests across 13 thresholds, all 78 possible combinations of two thresholds, and a LASSO-PGS of all 13 with bootstrapping 30 times. Although there were combinations of two PGS that performed better than the average LASSO-PGS, in practice it is challenging to anticipate which pairs of PGS will perform well. Thus, the LASSO, and possibly other machine learning approaches, provides an automated, robust way to combine PGS obtained from multiple P-value thresholds. In fact, when averaging across all single PGS models, the nomogram separation (i.e., difference between the top 30% and the bottom 30%) was about 82 mm3 (∼2% of average HV) and aligns with our previous work that used a single threshold PGS at P=0.75 [7]. In healthy older individuals with an estimated atrophy rate of about 1.4% [38] the difference is equivalent to 1.5 years of normal aging. The average nomogram separation for the LASSO-PGS models was about 118 mm3, which would be equivalent to 2.4 years of age-related atrophy (Figure 3).

Machine learning and precision diagnostic approaches using genetics and medical imaging share common challenges when being adopted in clinical settings. Perhaps the major challenge is to ensure that trained models and biomarkers perform well in the presence of a distribution shift of input values in the application setting compared to the development setting [39] and processing pipelines (e.g., different software versions). This same challenge applies to normative models, especially ones that combine multimodal data. Here such shifts can occur due to changes in technical parameters (e.g., MR scanner hardware, MR pulse sequence, or genotyping technology). We harmonized the SNP content prior to PGS generation to ensure the scores are comparable between datasets. We further harmonized the generated PGS and the imaging biomarkers using ComBat harmonization [40] to ensure comparable feature distributions. In our evaluations we observed a decline in performance (i.e., increase in prediction errors and a decrease in correlation) when transitioning from the UKBB test set, which shares the feature distributions with the training set (including same scanner), to the independent samples in ADNI and EPAD (Figure 4). However, this performance decline was driven by people with cognitive decline in ADNI. In fact, when evaluated separately, the performance on EPAD, which comprises participants at risk of dementia, is comparable to the UKBB test set (Suppl. Figure 9). Likewise, when grouping ADNI participants by their baseline diagnosis, the model performs even better than on the test set (Suppl. Figure 10), whereas prediction for participants with MCI and dementia are worse. The improved performance in CN participants in ADNI compared to the UKBB test set is likely due to better characterisation of cognitive decline in ADNI compared to UKBB overall, which may contain several participants with undiagnosed cognitive disorders. Thus, our results underline that the proposed harmonization steps produced models that can reliably be applied to novel data sets. Moreover, the reduced performance in people with diagnosed cognitive decline exemplifies that the concept of the normative model works as intended.

Previous studies have demonstrated that hippocampal volume is a relevant biomarker for disease progression in Alzheimer’s disease [41, 42]. Indeed, change in hippocampal volume is used as a primary and secondary endpoint in clinical trials [43–46] and correlates well with several neurocognitive scores [47–49]. In the same vein, estimates from normative models using hippocampal volume exhibit a strong association with clinical diagnosis and cognitive performance [3, 50]. Our results have demonstrated that enhancing normative models with genetic information improved association with neurocognitive test performance. In particular, we quantified the change in assigned percentile by a normative model that uses PGS (ASIP-LASSO) compared to the percentile assigned by a classic normative model that uses only demographic information (e.g., ASI model); a value we termed *model shift*.

Analysing the effect of *model shift* on the neurocognitive scores enabled us to quantify the added value of incorporating genetic information directly into the normative model. In the two assessed out-of-sample cohorts *model shift* was significantly associated with MMSE and CDR-SB both cross sectionally and longitudinally, when comparing to normative models without genetics (ASI model) or models incorporating single threshold PGS.

Importantly, we find that the association was not driven by few subjects that demonstrated substantial *model shifts* (i.e., >10%). Thus, even subtle adjustment of the assigned percentile improved the association with neurocognitive performance.

Notably, the magnitude of the *model shift* did not show any bias regarding diagnostic groups in ADNI (Supp. Figure 15). However, participants with dementia were more likely than others to have model shift values close to zero. This results from the fact that, in the dementia group, disease effects have reduced the HV substantially. Thus, participants receive very low percentile scores in the basic normative model (e.g., 2%), which remain unchanged after genetic adjustment. When test sub-scores were grouped by cognitive domain, we found memory and executive function to be the most significant cross-sectionally, both are domains with well-established links to the hippocampus [51–55].

*Model shift* did not only influence the baseline cognitive scores but also the participants longitudinal trajectory. In particular, the genetically adjusted normative model improved the prediction of longitudinal decline of MMSE and CDR-SB in ADNI (Table 2). More precisely, MMSE in ADNI (Supp. Table 3) demonstrated a *model shift* coefficient of .03, and a model-shift-by-time interaction coefficient of .02. The positive model-shift-by-time interaction coefficient (.02) represents that that the *model shift* effect leads to an increase over time (or rather a net slower decline when all variables are considered). Longitudinally, the orientation domain, which is specifically linked to the hippocampus [56, 57], showed the strongest influence by *model shift*. In ADNI, we observed that falling below the 10^th^ percentile using the genetics improved normative model results in a difference in CDR-SB (ADAS13) of 0.57 (1.74) over 18 months, which is comparable to the effect of Lecanemab in the clinical trial over the same period [58].

We have introduced a normative model that integrates neuroimaging biomarkers and genetics in the form of multiple PGS combined using the LASSO and evaluated its performance in two independent out-of-sample datasets. One major limitation is the level of data harmonization across these datasets: the three datasets used different MR hardware, MR pulse sequences, different versions of the processing software and genotyping arrays. We used readily available and widely used techniques (such as imputation [59] and NeuroComBat harmonisation [40]) to harmonise the data *post hoc*. In an ideal scenario, parameters linked to data acquisition and processing should be as closely matched as possible to maximise model performance and minimise confounding variance. However, the post hoc harmonisation applied here resembles real world applications more closely. A second limitation is that our normative model focused on assessing the size of the hippocampus. This selection was driven by the prominent role of the hippocampus for disorders such as Alzheimer’s disease, Depression, Schizophrenia, and others [60]. However, due to its shape and location the assessment of HV is fraught with uncertainties. While test-retest reliability has improved across recent Freesurfer versions [61], the hippocampus remains among the least reliable subcortical structures for volume estimation [62]. Thus, relying on the standard FreeSurfer pipeline may not be ideal and dedicated pipelines focusing on the hippocampus may result in more accurate estimates of *in vivo* hippocampal volume. Nevertheless, with this standard pipeline we demonstrated a significant improvement by incorporating genetic information. Moreover, more accurate HV estimates will percolate through the modelling pipeline and improve the model’s performance overall. Lastly, there are limitations regarding the training and evaluation data. We used UKBB as training cohort, where the disease phenotyping is incomplete and thus the dataset may contain undiagnosed participants. Moreover, the age range of participants in UKBB does not extend beyond 82 years, thus, older participants in ADNI and EPAD could not be evaluated. Future models should include further datasets with older participants.

In conclusion, we have demonstrated that polygenic scores for the same phenotype at different thresholds contain complementary information, which can be leveraged to build more predictive PGS of HV. We have further integrated these genetic predispositions for HV into normative models and shown that these enhanced models improve the prediction of neurocognitive test performance in two out-of-sample datasets. This study showcases how medical imaging data and genetics can be effectively combined to build tools that enable precision medicine. Our analysis focused on HV and dementia, but we believe this method has potential to improve a wide range of PGS applications for areas beyond neuroimaging.

## 4. Methods

### 4.1 PRE-PROCESSING

In total, three datasets were used in this study, summarized in Table 1.

#### 4.1.1 UKBB

The UK Biobank (UKBB) is a population cohort study of around 500k participants from the UK and contains a vast wealth of data including imaging, genetics, and clinical biomarkers [31]. From a total of 501,172 (272,747 female) participants (application number 65299), we excluded participants with no imaging or genetic data. To ensure HV models represented the spectrum of healthy aging, participants were excluded based on history of impairment (neurological or psychiatric disorders, head trauma, substance abuse, or cardiovascular disorders). And finally, to control for population level genetic heterogeneity, only participants with self-reported ‘British’ ethnic backgrounds were considered. Imaging and genetic protocols are described in [31] and [63], respectively. The dataset preparation followed the process described by [64], with HVs calculated using FreeSurfer V6 [65].

#### 4.1.2 ADNI

The Alzheimer’s Disease Neuroimaging Initiative (ADNI) was launched in 2003 as a public-private partnership, led by Principal Investigator Michael W. Weiner, MD. The primary goal of ADNI has been to test whether serial magnetic resonance imaging (MRI), positron emission tomography (PET), other biological markers, and clinical and neuropsychological assessment can be combined to measure the progression of mild cognitive impairment (MCI) and early Alzheimer’s disease (AD). Genetic and Imaging protocols are described in [66, 67] respectively. From a total of 2,430 (1,157 female) participants available in the ADNI dataset (downloaded Sep 2024), aged 50-91, we included the subjects with Freesurfer V7 imaging/HV available as described in [68]. Furthermore, we used genotyping data of ADNI participants pre-processed as previously described by (Scelsi et al., 2018) [69]. We filtered out subjects with < 0.8 predicted European ancestry based on SNPWeights Genetic Principal Components (GPCs) [70].

#### 4.1.3 EPAD

The European Prevention of Alzheimer’s Disease study (EPAD) is a growing longitudinal study of over 2500 (mostly non-demented) participants over the age of 50 with imaging, genetic, and clinical data, collected from 31 sites across Europe (Ritchie et al., 2019) [33]. Data from a total of 2660 subjects (1457 female) aged 50 – 89 from were obtained with HVs calculated using Freesurfer V6. We filtered out subjects with < 0.8 predicted European ancestry based on SNPWeights Genetic Principal Components (GPCs).

### 4.2 Polygenic Scores

We built our Polygenic Scores using Clumping and Thresholding (C+T) with PRSice V2 [34]. We used a previously reported HV GWAS of 26,814 European subjects from the ENIGMA study [71]. We filtered for minor allele frequency of 0.05, genotype missingness of 0.1, and performed clumping at 250Kb. We calculated PGSs at 13 P-value thresholds (1E-8, 1E-6, 1E-5, 1E-4, 1E-3, 0.01, 0.05, 0.1, 0.2, 0.4, 0.5, 0.75, 1) using the sum of effect sizes weighted by SNP allele count. Next, we converted the PGSs to Z-scores and regressed out the effect of genetic principal components (GPCs) using linear regression. We used 40 GPCs previously calculated from a select independent set of SNPs from UKBB subjects using fastPCA [72]. To compute the double PGS, i.e., a PGS considering polygenic scores at two P-value cutoffs, their corresponding Z-scores are summed.

#### 4.2.1. LASSO-PGS

We derived a LASSO-PGS by using weights from a LASSO regression to linearly combine the PGSs at all 13 thresholds after Z-scoring. Here the target variable in UKBB was bilateral HV and the inputs were the 13 PGS at different P-value thresholds. We used k-fold cross-validation to tune the regularization parameter (λ). Across all experiments, LASSO weights were derived from a UKBB training set and applied to the testing sets (UKBB testing, ADNI and EPAD). To achieve robust statistics, LASSO training was bootstrapped 30 times.

#### 4.2.2 Cross Dataset Common Polygenic Score

For experiments that were carried out across datasets, it was required to harmonize the PGS. First, we only considered SNPs that were available in all three databases for the PGS computation (genotyped UKBB, imputed ADNI and EPAD). Next, we estimated genetic ancestry using the SNPWeights tool [70], which is based on a large external reference panel. We used NeuroCombat [40, 73] to harmonize the PGS values to UKBB by retaining the effect of the SNPWeights GPCs. After the harmonization step, we linear regressed out the effect of the GPCs on PGSs across all three datasets.

### 4.3. Hippocampal Volume

For HV and ICV, we computed bilateral HVs from left and right HVs generated by FreeSurfer [65] and we used estimated total intracranial volume (eTIV) as our ICV measurement. HVs were generated using FreeSurfer v6 for EPAD and UKBB and FreeSurfer v7 for ADNI.

#### 4.3.1. Cross-dataset Harmonization

Similarly to the harmonization of the PRS values, also the imaging biomarkers must be harmonized to account for biases induced by study protocols and processing pipelines. We used NeuroCombat [40, 73] to harmonize the HV and eTIV values to the UKBB. This was conducted in two steps: first eTIV was harmonized retaining the effects of age, sex and GPCs. Second, we harmonized the HVs retaining the effects of age, sex, diagnosis, harmonized eTIV and PGS.

#### 4.3.2 PGS Correlations

We computed the Pearson’s correlations of the 13 scores with each other and with HV. Furthermore, we investigated the ranking of subjects by PGS in two ways. Firstly, we directly examined how the PGS-based ordering of subjects changed across PGSs. Secondly, we computed the overlap between the top/bottom 30% of subjects sorted by each of the 13 PGSs.

### 4.4 Estimating Nomogram Separation

For each PGS (single/double/LASSO), we build nomograms from the top vs bottom 30% scoring participants using Gaussian Process Regression (GPR). The method follows our previously described work [7]. Briefly, our GPR models were trained with the laGP [74] R library, with the commonly used squared exponential covariance kernel function. After training, we build nomograms by predicting HVs within the training age range with increments of 0.01 years and calculating the quantile curves at nine percentiles: 2.5%, 5%, 10%, 25%, 50%, 75%, 90%, 95%, and 97.5%.

To derive the difference between two nomograms, we computed the average difference between their corresponding quantile curves. To this end, we first restricted the range of the nomograms to the intersection of their training data age range, because of GPRs tend to diverge quickly where no training data is present rendering comparisons outside this range unreliable. Next, we calculated the difference of two nomograms across the age range at their matching percentile curves (e.g., at 2.5%). The nomogram difference is then defined as the average difference between all pairs of quantile curves.

### 4.5 Multivariable Normative Model

#### 4.5.1 Model Training

In our previous work [7], stratification lead to nomogram separation when comparing top/bottom strata, but when the middle strata was included (top vs bottom 50%) nomogram separation is greatly diminished. Hence, we can make better use of the data by building higher dimensional GPR models instead. Higher dimensional models are capable of learning from all the input covariates (age, ICV, PGS, etc.) jointly and they can still produce nomograms for different strata of covariates. The extracted z-score is the key information to be extracted from normative models, which can be done with GPRs of any dimension.

We trained GPRs with 17 progressively complex sets of covariates. The models produce HV z-scores given only Age (A Model), and Sex (AS Model), and ICV (ASI Model), and PGS (ASIP Model), where PGS is either the LASSO-PGS (ASIP-LASSO) or one of 13 single threshold PGSs (ASIP-single: ASIP-1E-8 to ASIP-1). To achieve robust LASSO models, the UKBB was randomly split in half 30 times and both LASSO-PGS and ASIP-LASSO were generated each time. For model assessment, we report the root mean squared error (RMSE) and the Pearson’s correlation coefficient of the z-scores to HV in the training, testing, and out-of-sample tests.

Importantly, we limit our testing to subjects whose ages and ICVs are within the range seen by the trained model, since GPR diverges quickly when extrapolating outside the range of training values.

#### 4.5.2 Neurocognitive Tests

We tested the association between improved normative models in out-of-sample data on range of neurocognitive measures depending on their availability in the corresponding dataset. For example, both ADNI and EPAD had Mini-Mental-State-Examination (MMSE) and Clinical Dementia Rating Sum of Boxes (CDR-SB), while ADNI also provides the Alzheimer’s Disease Assessment Scale (ADAS), and EPAD had the Repeatable Battery for the Assessment of Neuropsychological Status (RBANS).

We report the significant p-values after Holm–Bonferroni correction across five tests in each dataset independently. We also looked at the performance across five domains (Memory, Language, Orientation, Visual-Spatial, Executive Function) by combining subtests of NC measures where available. For example, in ADNI, the Memory domain consisted of: Everyday Cognition – study partner memory score, MMSE recall task, ADAS memory task, and the MOCA memory task. The details of which tests were used from each database, and which sub-tests composed every domain can be found in Supp. Tables 2 and 3. Tests were conducted for each of the sub-tests independently and then combined using Stouffer’s method adjusted for dependent P-values (implemented in the *poolr* R library [75]) to provide a domain level P-value.

#### 4.5.3 Defining Model Shift

The aim is to quantify the improved association between NC values and GPR normative models constructed using the LASSO-PGS compared to classic normative models. In order to compare the z-scores from two GPR models, we consider the difference between them in terms of the subjects’ predicted percentiles. Specifically, we look at how much the percentile shifts when going from ASIP-LASSO to any other model (e.g., ASI), a value we term *model-shift*. If a subjects percentile in the ASIP-LASSO model is higher than in the other model, their model-shift would be positive, e.g., +5%, and vice versa. Thus, positive model-shifts suggest that the ASIP-LASSO model rates the HV value more normal. Thus, we expect positive model-shifts to represent better neuro cognitive function such as better memory.

#### 4.5.4 Cross Sectional Analysis

We used linear regression to assess the effect of model-shift (in percentiles) on the cognitive measures. These models were adjusted for standard covariates such as age, sex, ICV and the percentile from a GPR model, which ASIP-LASSO was compared to. We also conducted two sensitivity analyses: (i) to ensure that our results were not driven by subjects with high model-shifts, we performed the same tests while excluding subjects where model shift magnitude was greater than 10% and (ii) and in ADNI, to ensure our results were not driven completely by people with dementia, we also performed the tests after excluding participants diagnosed with Dementia at the baseline visit.

#### 4.5.4 Longitudinal Analysis

Leveraging the longitudinal design of the ADNI and EPAD studies, we used linear mixed effects models to assess whether the model-shift had longitudinal impact. More precisely, we estimated three sets of models. One with age, sex, ICV, baseline percentile, visit-time, baseline percentile and time interaction as fixed effects and subject ID and visit-time as random effects (termed ‘*MS-none*’ since model-shift is not included). A second model that adds model-shift as a fixed effect on top of MS-none (*MS-fixed*). and a third that adds the model-shift and time interaction as fixed effect on top of MS-fixed (*MS-time*). We then report the results of an ANOVA test between MS-none versus MS-fixed, and MS-fixed versus MS-time. We performed this test with ASIP-LASSO versus ASI model-shift, and ASIP-LASSO versus each of the 13 single threshold ASIP-PGS model-shift. In order to illustrate the longitudinal results, we plotted, across time, the predicted performance with the mixed effects models for an average participant with +10% model-shift versus −10% model shift.

Similar to the cross-sectional tests, we conducted sensitivity analyses by excluding outliers in terms of |model-shift| ≥ 10% and excluding Alzheimer’s Disease subjects from ADNI.

#### 4.4.4 Crossing Label Longitudinal Analysis

Finally, we used linear mixed effects models to assess whether crossing a specific percentile curve between the ASI and the ASIP-LASSO models had any longitudinal impact. For each percentile (1-100), we defined a percentile *crossing-label* depending on whether the model-shift from ASI to ASIP-LASSO did not cross that percentile (0), crossed above it (−1), or crossed below it (+1). We then constructed linear mixed effects models with age, sex, ICV, baseline Z-score, crossing-label, visit-time, crossing label and time interaction as fixed effects, and subject ID and visit-time as random effects. We report p-values that, in addition to being corrected for five NC tests and five domains in each dataset, are also corrected for the 100 percentiles tested using FDR Correction.

## 5. Code Availability

The scripts and code used in this study have been made publicly available and can be found at: https://github.com/Mo-Janahi/Multi-Threshold-PGS. All underlying data, and derived quantities, are available by application from the UK Biobank at http://www.ukbiobank.ac.uk, ADNI at http://adni.loni.usc.edu/data-samples/access-data/, and EPAD as https://ep-ad.org/index.php/open-source-data/. Summary statistics from the genome-wide association studies described in this paper is available from the NHGRI-EBI GWAS catalogue, study number GCST003961. URL: https://www.ebi.ac.uk/gwas/studies/

## Acknowledgements

Data used in preparation of this article were obtained from the Alzheimer’s Disease Neuroimaging Initiative (ADNI) database (adni.loni.ucla.edu). Data collection and sharing for this project was funded by the Alzheimer’s Disease Neuroimaging Initiative (ADNI) (National Institutes of Health Grant U01 AG024904) and DOD ADNI (Department of Defense award number W81XWH-12-2-0012). ADNI is funded by the National Institute on Aging, the National Institute of Biomedical Imaging and Bioengineering, and through generous contributions from the following: AbbVie, Alzheimer’s Association; Alzheimer’s Drug Discovery Foundation; Araclon Biotech; BioClinica, Inc.; Biogen; Bristol-Myers Squibb Company; CereSpir, Inc.; Cogstate; Eisai Inc.; Elan Pharmaceuticals, Inc.; Eli Lilly and Company; EuroImmun; F. Hoffmann-La Roche Ltd and its affiliated company Genentech, Inc.; Fujirebio; GE Healthcare; IXICO Ltd.; Janssen Alzheimer Immunotherapy Research & Development, LLC.; Johnson & Johnson Pharmaceutical Research & Development LLC.; Lumosity; Lundbeck; Merck & Co., Inc.; Meso Scale Diagnostics, LLC.; NeuroRx Research; Neurotrack Technologies; Novartis Pharmaceuticals Corporation; Pfizer Inc.; Piramal Imaging; Servier; Takeda Pharmaceutical Company; and Transition Therapeutics. The Canadian Institutes of Health Research is providing funds to support ADNI clinical sites in Canada. Private sector contributions are facilitated by the Foundation for the National Institutes of Health (www.fnih.org). The grantee organization is the Northern California Institute for Research and Education, and the study is coordinated by the Alzheimer’s Therapeutic Research Institute at the University of Southern California. ADNI data are disseminated by the Laboratory for Neuro Imaging at the University of Southern California. As such, the investigators within the ADNI contributed to the design and implementation of ADNI and/or provided data but did not participate in analysis or writing of this report.

Data used in preparation of this article were obtained from European Prevention of Alzheimer’s Dementia Longitudinal Cohort Study (EPAD LCS). The authors express their most sincere gratitude to the EPAD LCS participants, without whom this research would have not been possible. EPAD is supported by the EU/EFPIA Innovative Medicines Initiative (IMI) grant agreement 115736. The project leading to this paper has received funding from the Innovative Medicines Initiative 2 Joint Undertaking under grant agreement No 115952. This Joint Undertaking receives the support from the European Union’s Horizon 2020 research and innovation programme and EFPIA. This communication reflects the views of the authors, and neither IMI nor the European Union and EFPIA are liable for any use that may be made of the information contained herein.

JMS is a National Institute for Health Research (NIHR) Senior Investigator and acknowledges the support of the NIHR University College London Hospitals Biomedical Research Centre and the UCL Centre of Research Excellence, an initiative funded by British Heart Foundation (RE/24/130013). He has grant funding from Alzheimer’s Research UK, LifeArc, Brain Research UK, Weston Brain Institute, Medical Research Council, British Heart Foundation, Wolfson Foundation, UK Dementia Research Institute and Alzheimer’s Association.

