## Supplementary Figures for "Incorporating Multi-threshold Polygenic Risk in Hippocampal-based Normative Models Improves Cognitive Decline Prediction"

Integrating Multiple Polygenic Scores with Normative Models Improves Cognitive Decline Detection in At-Risk Individuals

### 1. Supplementary Figures / Tables

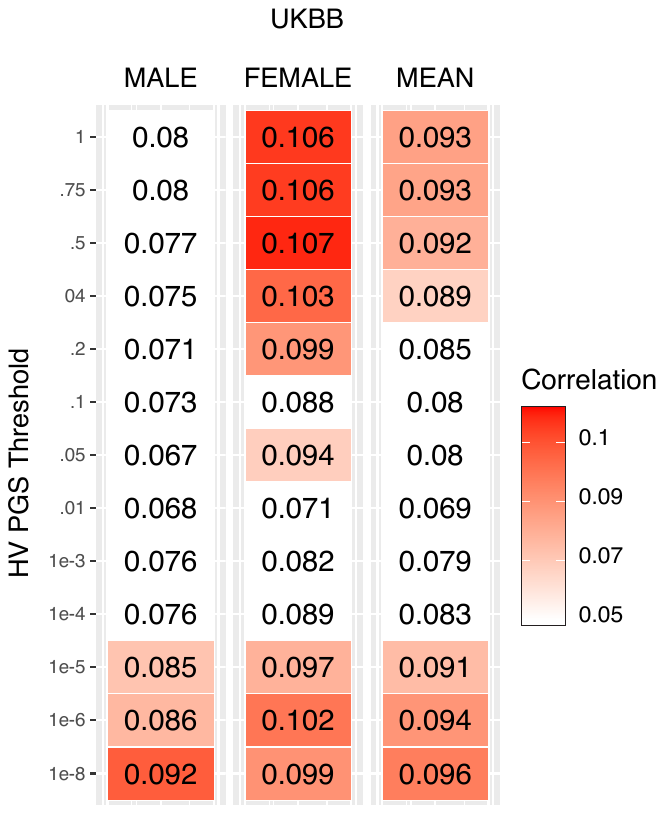

**Supplementary Fig. 1: Correlations of PGSs to Hippocampal Volume (HV) across UKBB strata.**

Pearsons Correlations (r) of bilateral HV in male/female UKBB participants with PGSs of bilateral HV generated at 13 different p-value thresholds. PGS correlate similarly to HV at opposing ends of the threshold spectrum.

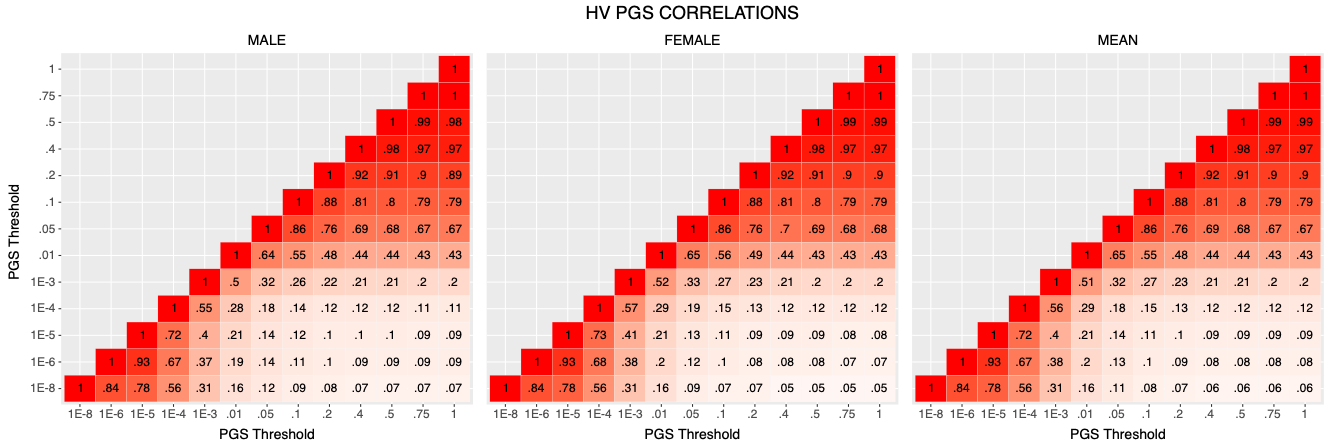

**Supplementary Fig. 2: inter PGS correlations across male/female strata.**

Pearsons Correlation (r) of bilateral HV PGS generated at 13 p-value thresholds to each other across UKBB male/female strata. Correlation between scores is directly correlated to the distance between the thresholds used to generate them.

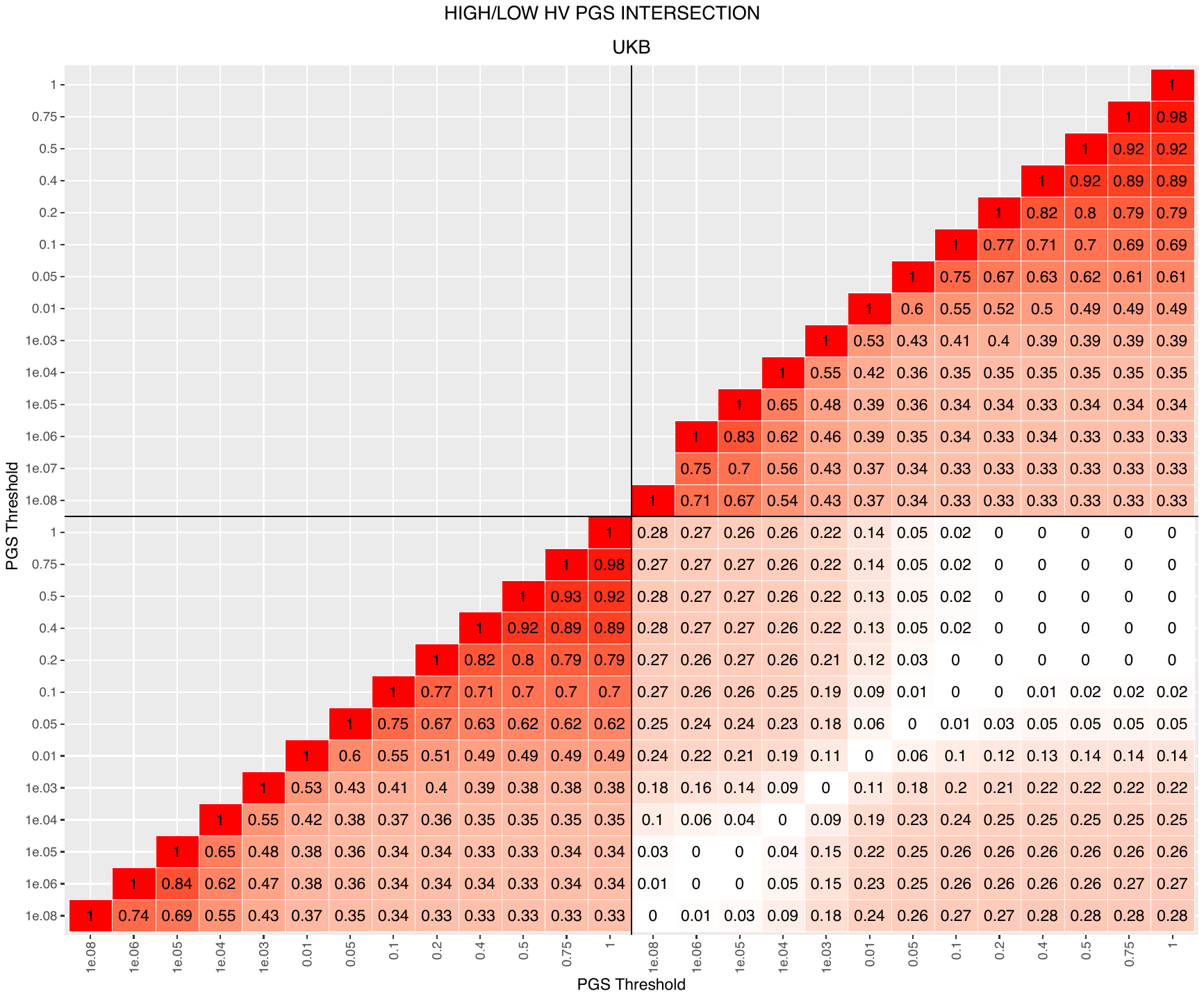

**Supplementary Fig. 3: Intersections of subjects across PGS thresholds.**

Percent of overlapping participants across top/bottom 30% bins of PGSs generated at 13 p-value thresholds. Top/top and bottom/bottom bins generally intersect better than top/bottom bins, but there is a large overlap between participants ranked in the highest bin in one threshold and lowest in another.

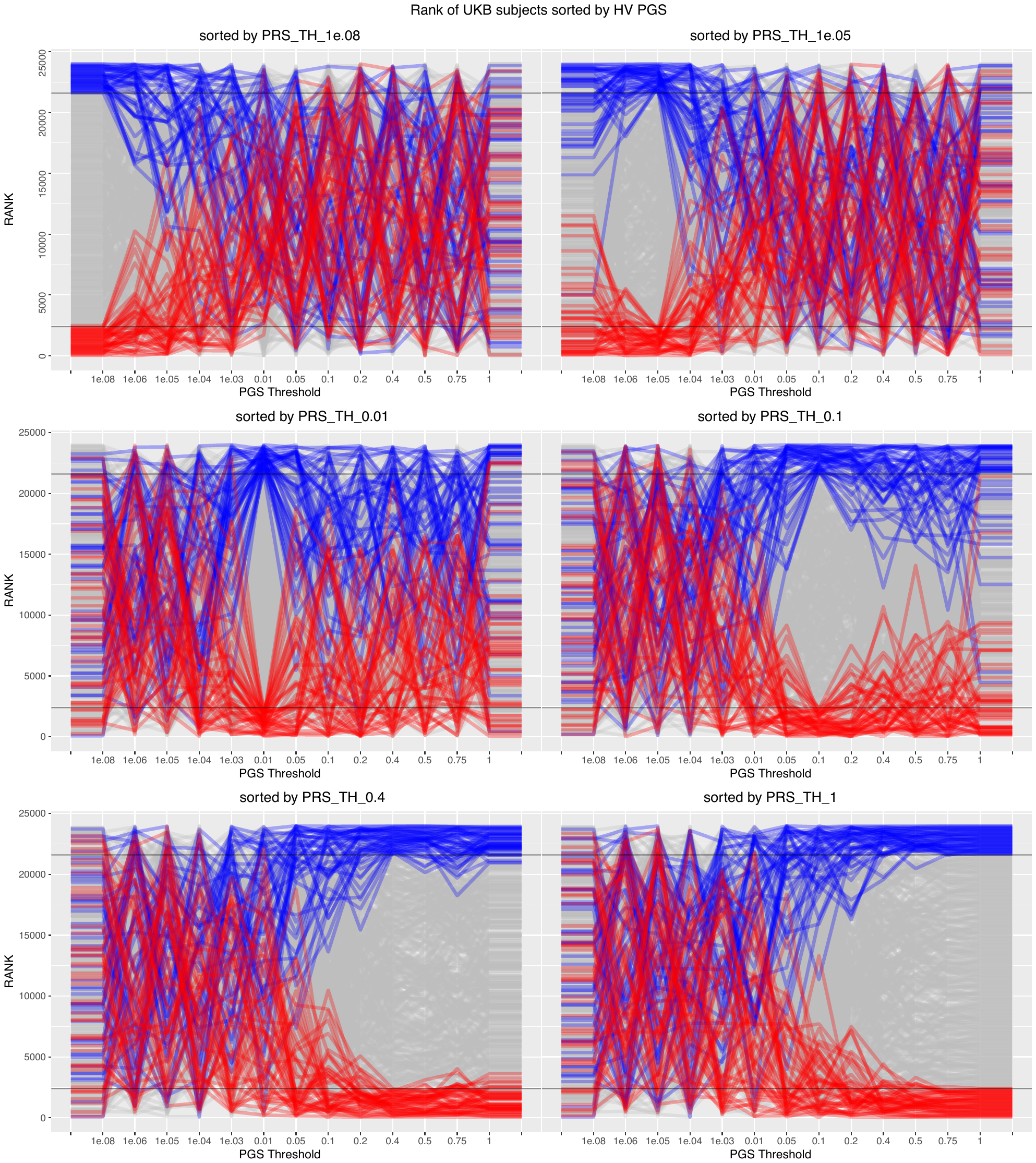

**Supplementary Fig. 4: ranking of subjects across PGS thresholds.**

The order of participants is not preserved across PGS thresholds. When ranking participants by the one threshold it is the exception that that ranking is maintained in another threshold. The top and bottom 10% of participants (blue/red lines) appear randomly distributed among the ranks of participants by distant threshold PGS.

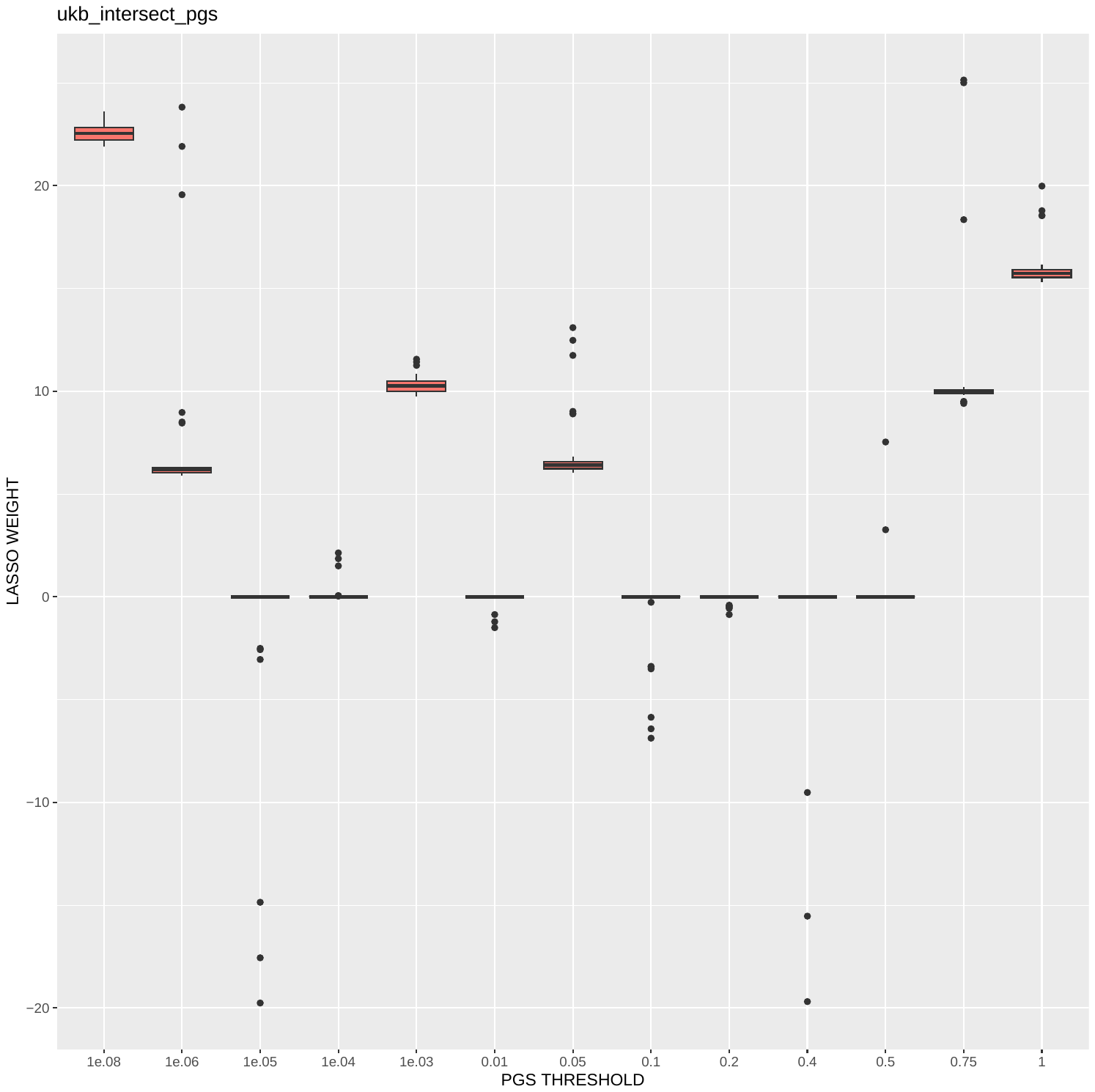

**Supplementary Fig 5: Average LASSO-PGS weights.**

We split UKBB participants in half randomly 30 times and trained a LASSO-PGS each time. Some thresholds are consistently chosen over others (P=1E-8 and P=1). For this figure, the PGSs were Z-scored before LASSO training to render the resulting weights comparable.

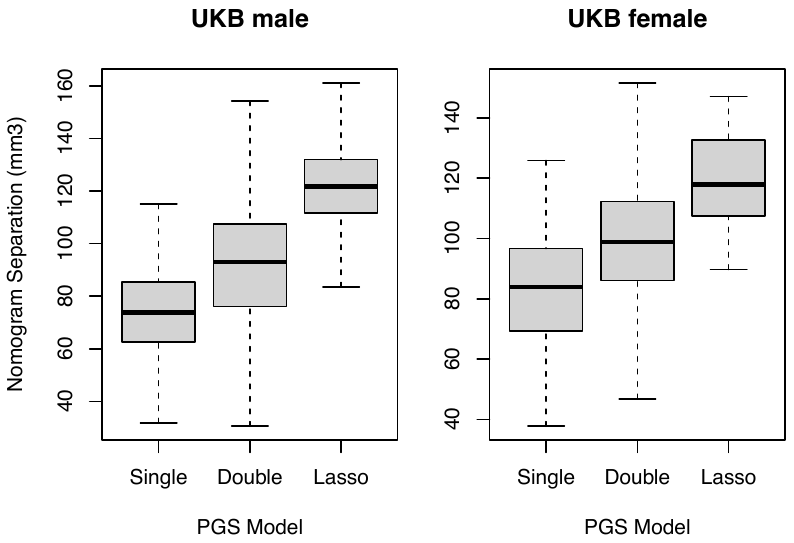

**Supplementary Fig. 6: Nomogram Separation across male and female strata.**

Comparing the nomogram separation in UKBB across male/female strata. Across 30 bootstraps of stratification by 13 single threshold PGSs; 78 combinations of two thresholds; and a LASSO of all thresholds, on average, double-PGS is 22% better than a (classic) single threshold PGS, and a LASSO-PGS is 50% better.

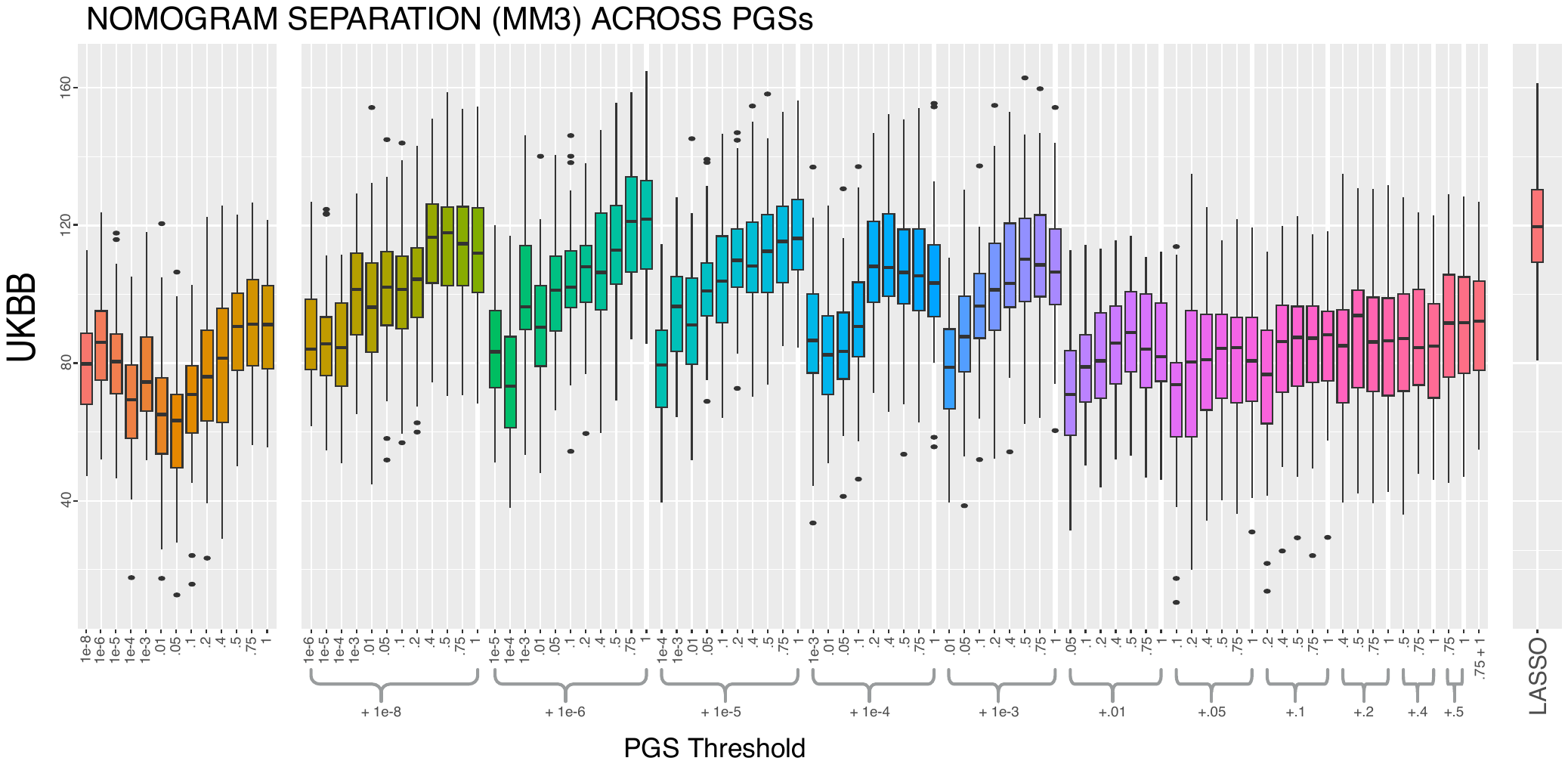

**Supplementary Fig. 7: Nomogram Separation across single/double/lasso threshold PGS.**

Showing, in detail, the nomogram separation across the 30 bootstraps of stratification by 13 single threshold PGSs; 78 combinations of two thresholds; and a lasso of all thresholds.

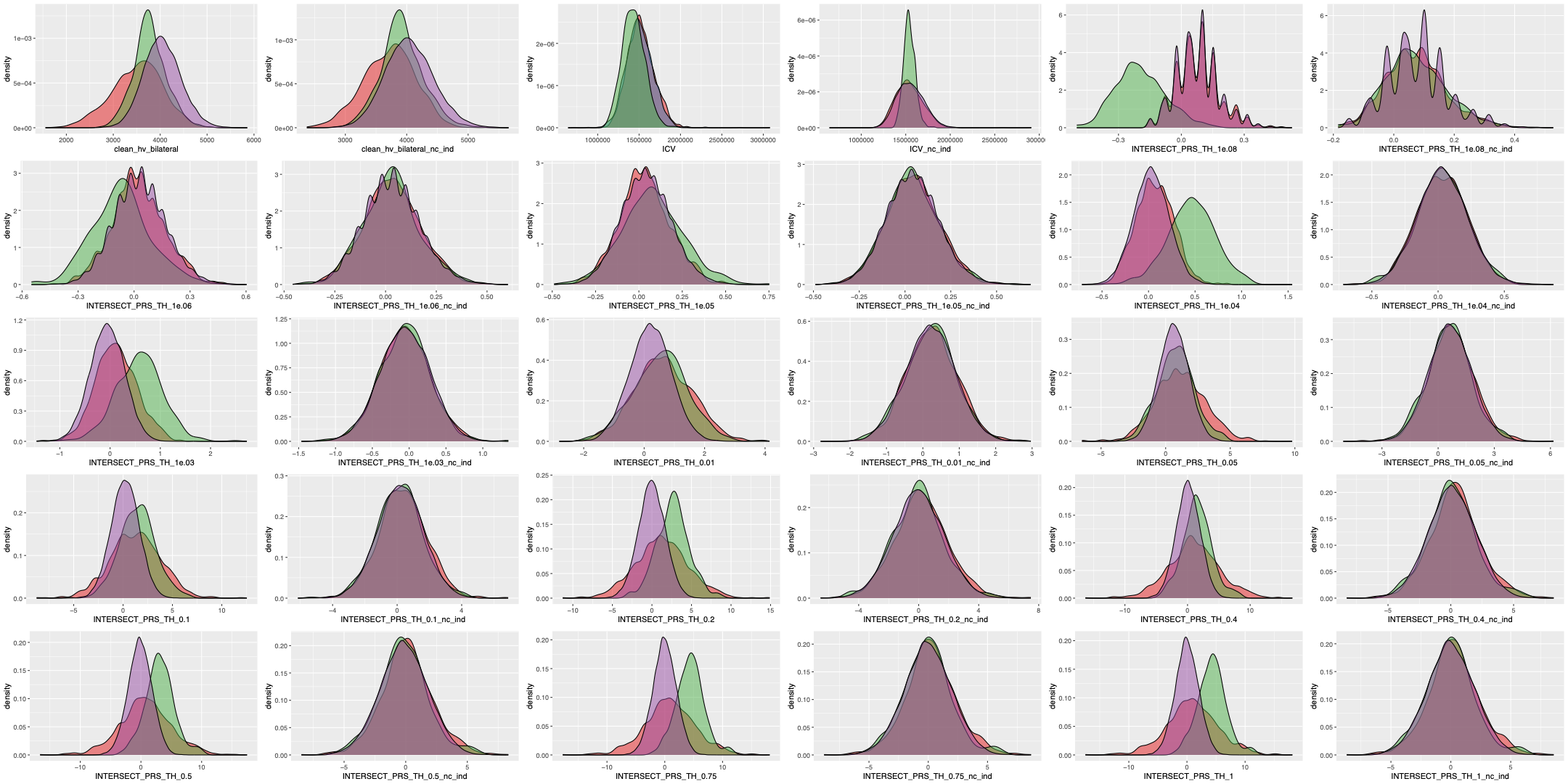

**Supplementary Figure 8: Distributions of harmonized and un-harmonized covariates across UKBB (Purple), ADNI (Red) and EPAD (Green).** Every column of un-harmonized covariates (1,3,5) is followed by its corresponding column of harmonized covariates (2,4,6).

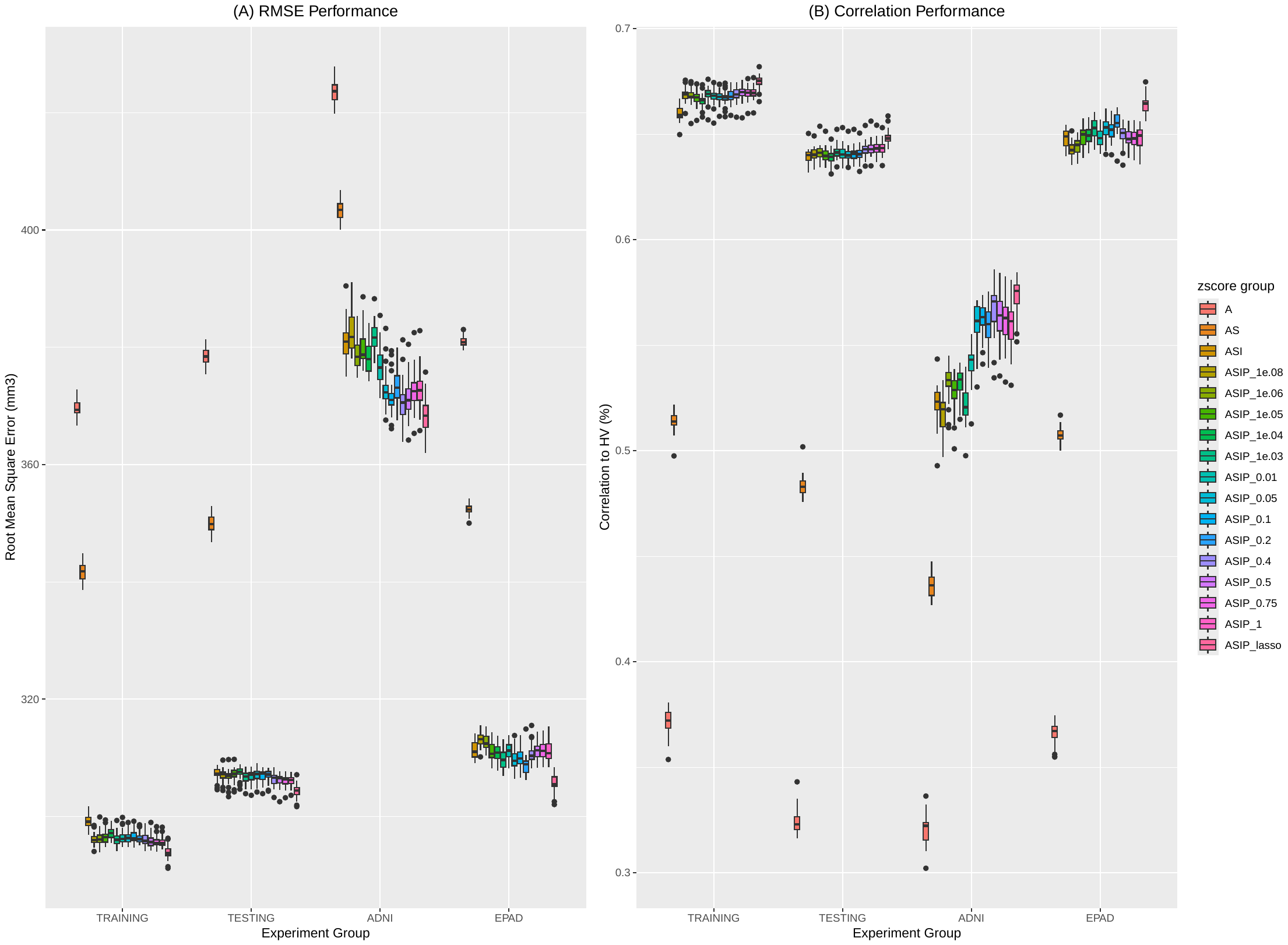

**Supplementary Fig. 9: GPR HV model assessment across all sets of covariates.**

(A) Root Mean Squared Error (RMSE) between predicted HV by the GPR models and measured HV. (B) Pearson’s correlation coefficient between predicted and measured HV across models. On average, the RMSE lowers when adding more covariates, and most when adding the LASSO score. RMSE increases and HV correlation decreases when going from training data to test data to out-of-sample cohorts. Much more so in ADNI compared to EPAD.

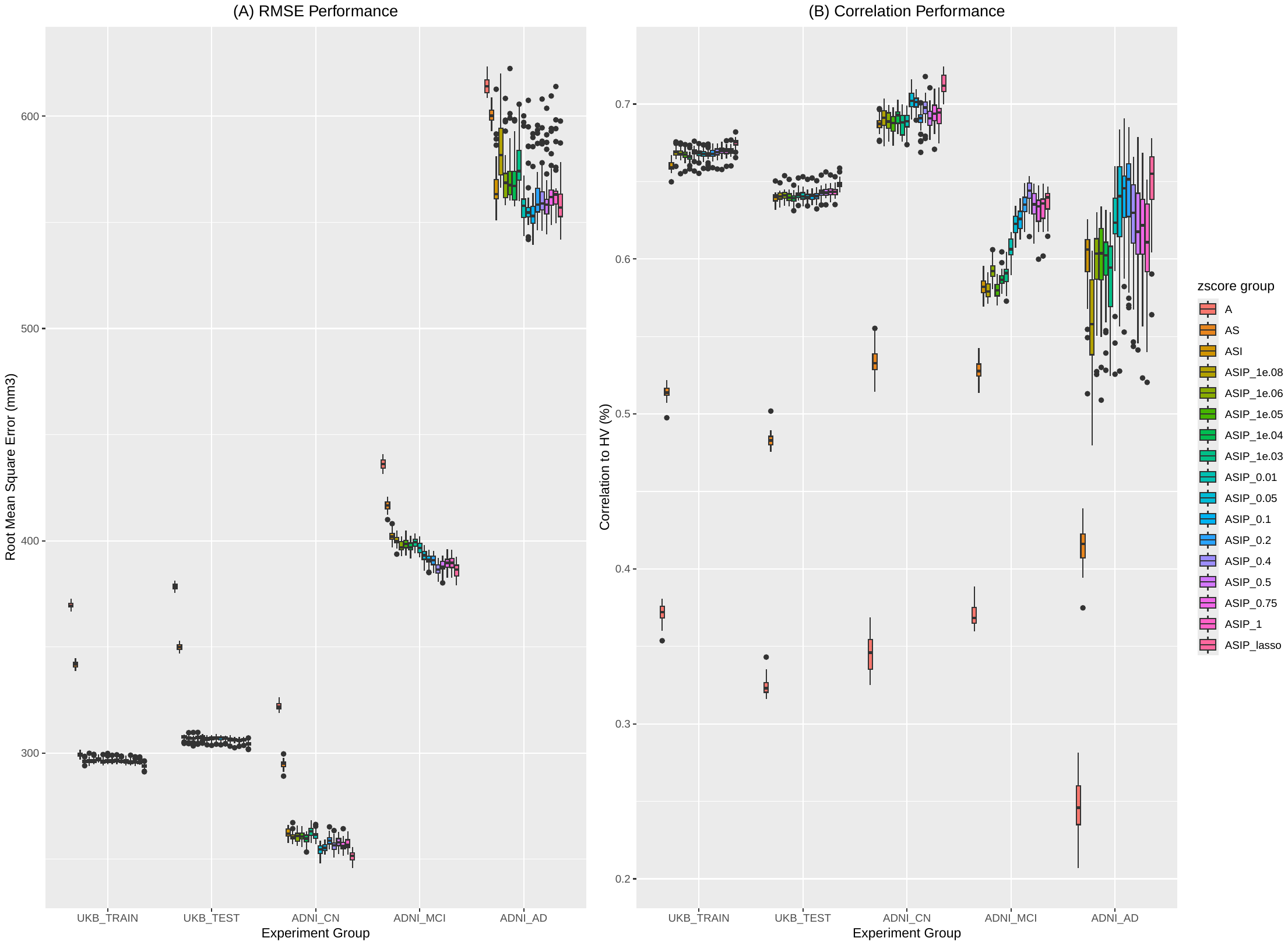

**Supplementary Fig. 10: GPR HV model assessment across all sets of covariates and diagnostic groups in ADNI.**

Root Mean Squared Error (RMSE) and Pearson’s correlation between predicted and measured HV across UKBB training/testing and ADNI diagnostic groups. Within diagnostic groups, RMSE lowers, and correlation increases when adding more covariates, and most when adding the LASSO score. CN ADNI subjects have lower RMSE and higher correlation than even the UKBB training set.

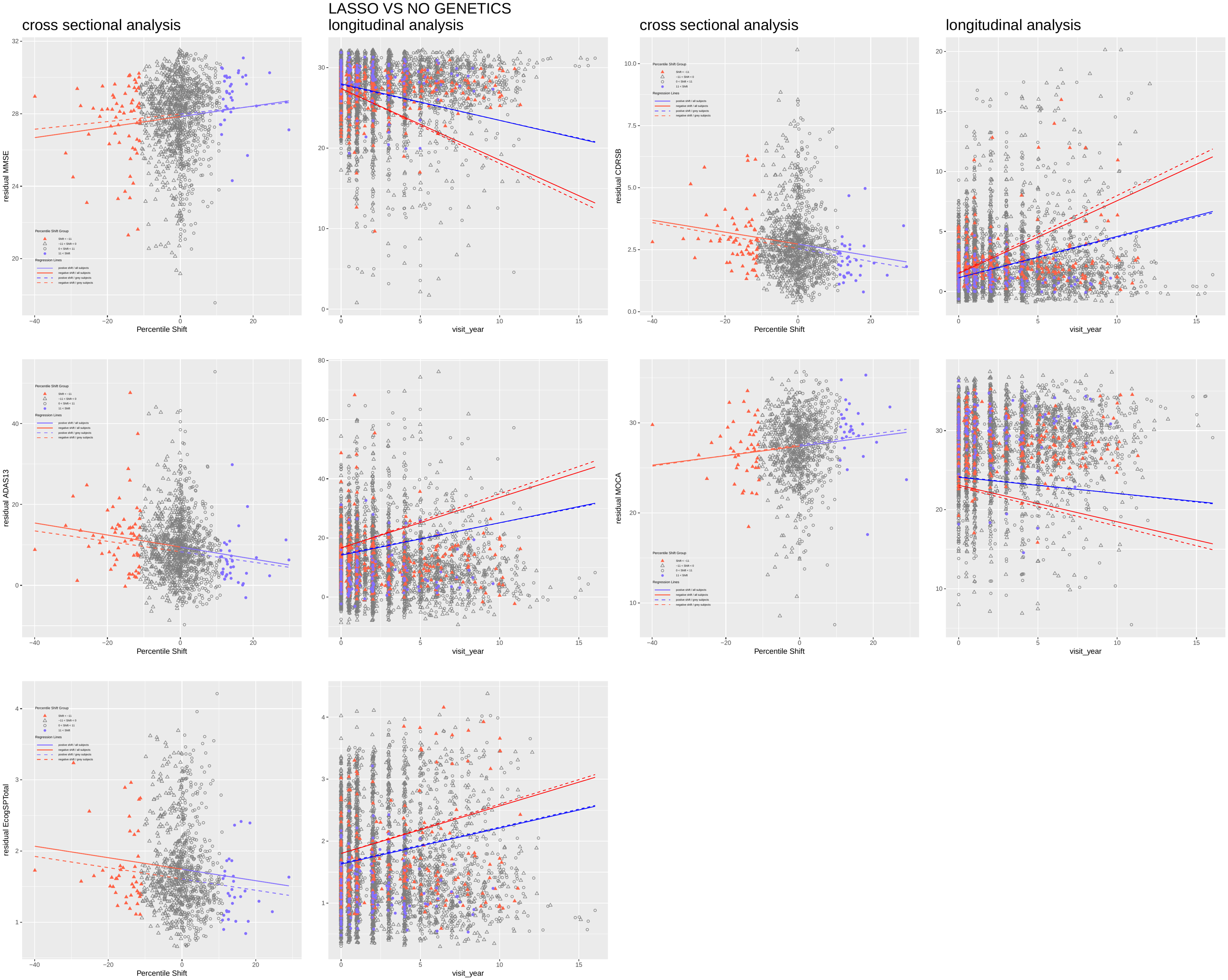

**Supplementary Fig. 11: ASI to ASIP-lasso model shift ADNI performance**

Comparing performance in MMSE, CDR, ADAS, MOCA, and ECOG in ADNI to ASIP-LASSO to ASI model shift. At baseline, model shift is positively correlated to tests where a higher score is better and visa-versa. Some correlations remain significant when only observing moderately shifted participants (|model-shift|<11%; grey points) and are not significantly changed (dashed line). These correlations remain significant longitudinally in linear mixed effects models. The regression lines drawn are projections/prediction for an average subject with +10 vs -10 model-shift (blue vs red).

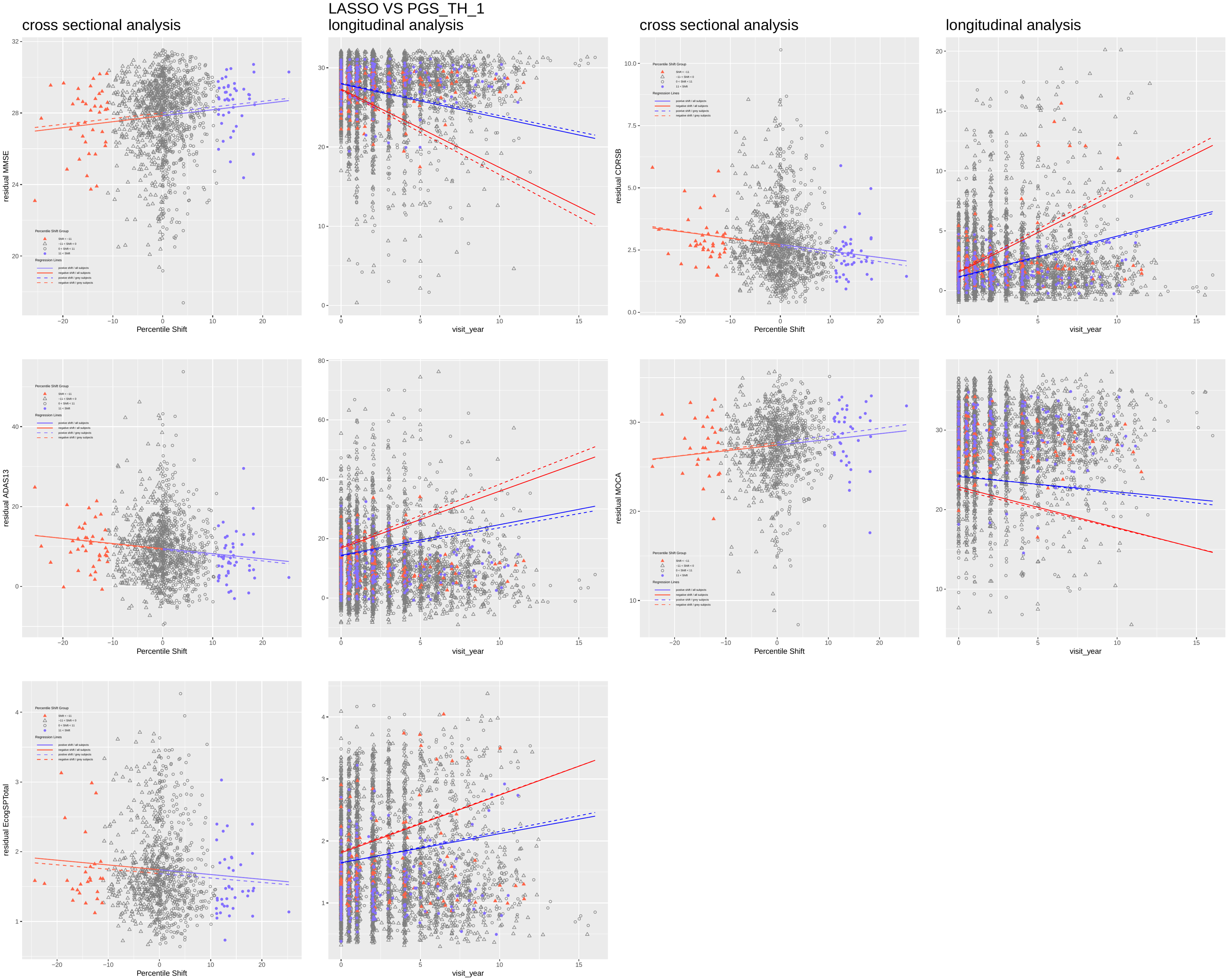

**Supplementary Fig. 12: ASIP-lasso vs ASIP-1 model shift performance in cross-sectional and longitudinal tests in ADNI.**

Comparing performance in MMSE, CDR, ADAS, MOCA, and ECOG in ADNI to ASIP-LASSO to ASIP-1 model shift. Results follow same lines as previous figure but less pronounced overall.

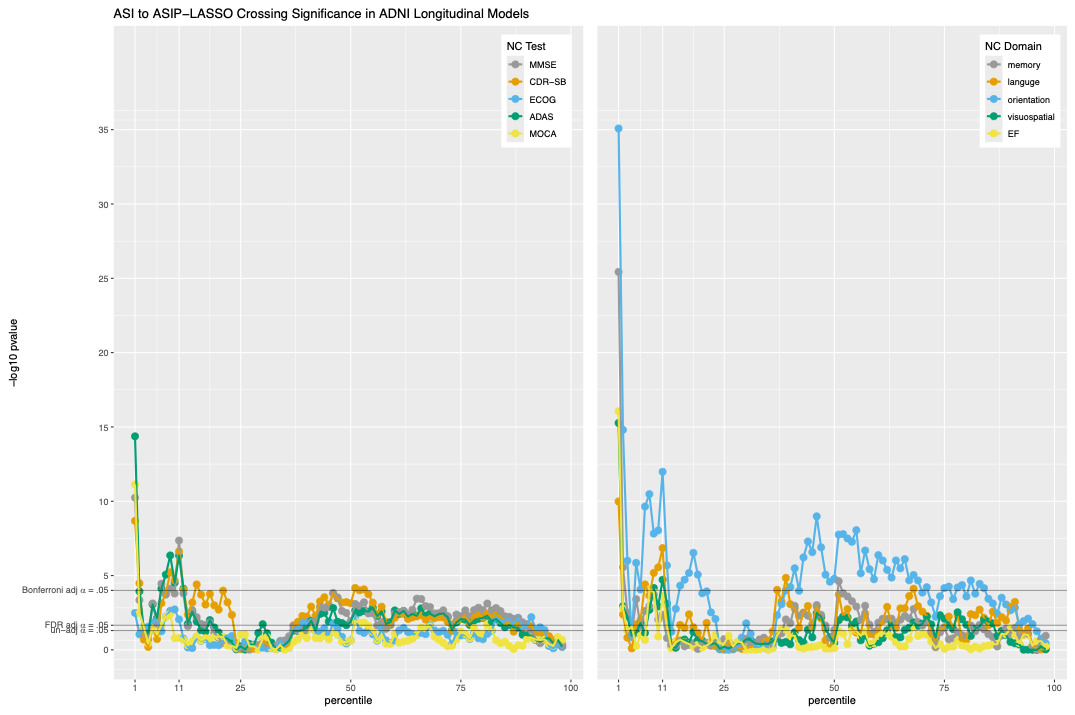

**Supplementary Fig. 13: Crossing Label P-Value.**

Labelling if ASI to ASIP-LASSO model shift crosses specific percentile thresholds reveals that some thresholds are more significant than others. In ADNI, model-shift crossing the 1^st^ percentile is significantly correlated to five neurocognitive tests and all five neurocognitive domains even after Bonferroni adjustment for 500 tests (5 NC test or domain * 100 percentiles). Similar, yet less significant results are observed at the 11^th^ and 50^th^ percentile. Horizontal lines depict 95% significance levels for uncorrected, FDR corrected, and Bonferroni corrected levels.

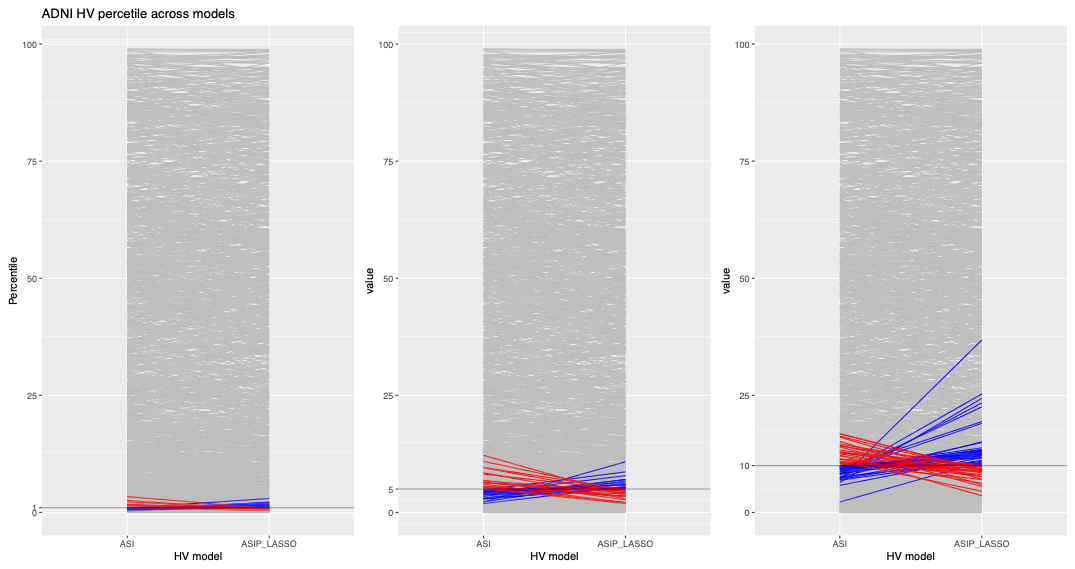

**Supplementary Fig. 14: ASI to ASIP-LASSO model-shift across percentiles.**

Illustrative examples of ADNI subjects with scores from ASI and ASIP-LASSO models. For 3 selected percentiles (1%, 5%, 10%), subjects whose model-shift crosses the percentile are highted (blue crossing above, red crossing below).

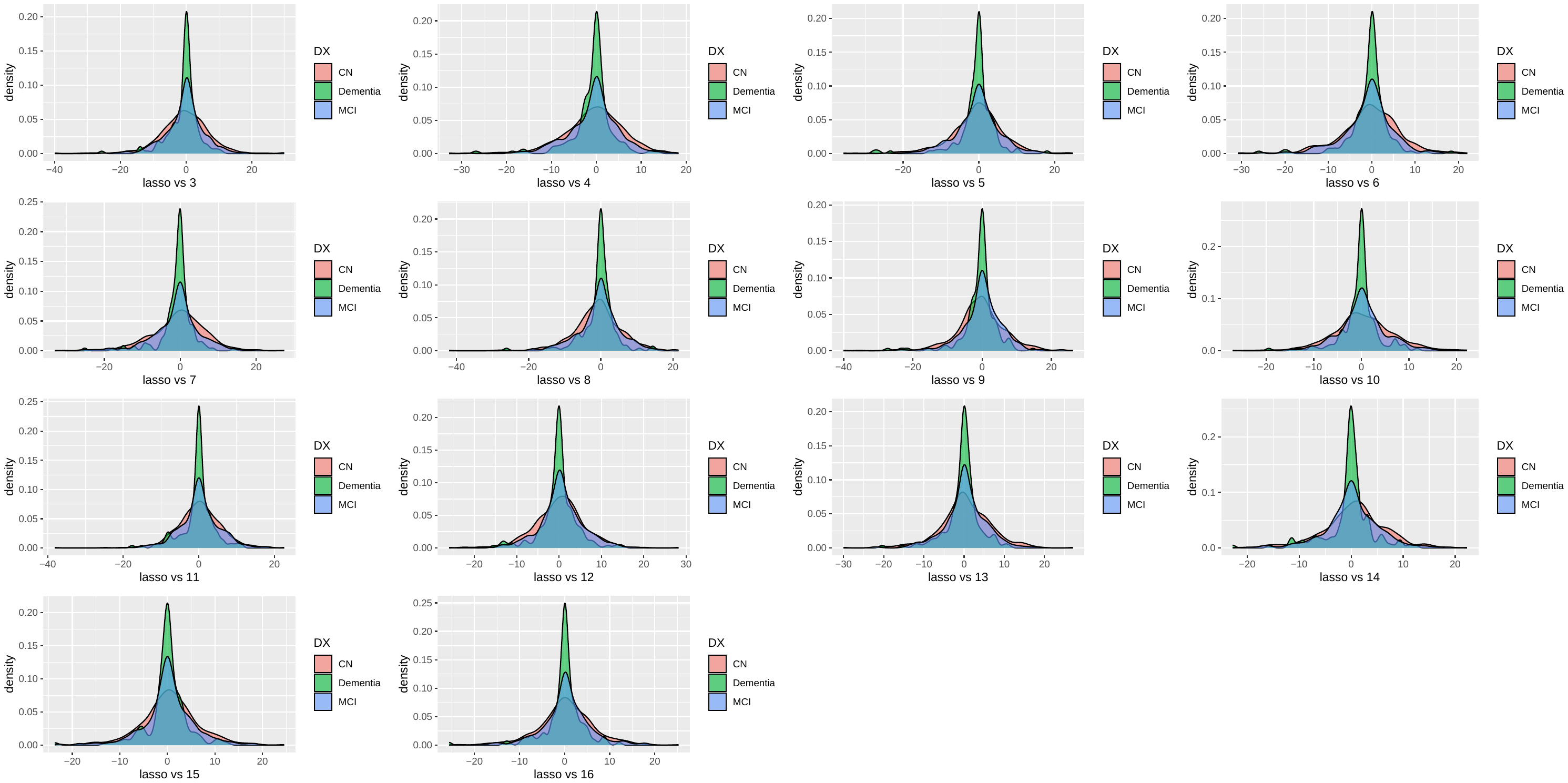

**Supplementary Fig. 15: Model shift across ADNI diagnoses strata.**

Model shift histograms grouped by ADNI diagnostic strata. No bias is seen in any diagnostic group for any model-shift.
